# Hierarchical and fine-scale mechanisms of binocular rivalry for conscious perception

**DOI:** 10.1101/2023.02.11.528110

**Authors:** Chencan Qian, Zhiqiang Chen, Gilles de Hollander, Tomas Knapen, Zihao Zhang, Sheng He, Peng Zhang

## Abstract

Conscious perception alternates between the two eyes’ images during binocular rivalry. How hierarchical processes in our brain interact to resolve visual competition to generate conscious perception remains unclear. Here we investigated the mesoscale neural circuitry for binocular rivalry in human cortical and subcortical areas using high-resolution functional MRI at 7 Tesla. Eye-specific response modulation in binocular rivalry was strongest in the superficial layers of V1 ocular dominance columns (ODCs), and more synchronized in the superficial and deep layers. The intraparietal sulcus (IPS) generated stronger eye-specific response modulation and increased effective connectivity to the early visual cortex during binocular rivalry compared to monocular “replay” simulations. Although there was no evidence of eye-specific rivalry modulation in the lateral geniculate nucleus (LGN) of the thalamus, strong perceptual rivalry modulation can be found in its parvocellular (P) subdivision. Finally, IPS and ventral pulvinar showed robust perceptual rivalry modulation and increased connectivity to the early visual cortex. These findings demonstrate that local interocular competition arises from lateral mutual inhibition between V1 ODCs, and feedback signals from IPS to visual cortex and visual thalamus further synchronize and resolve visual competition to generate conscious perception.

**Graphical abstract:** 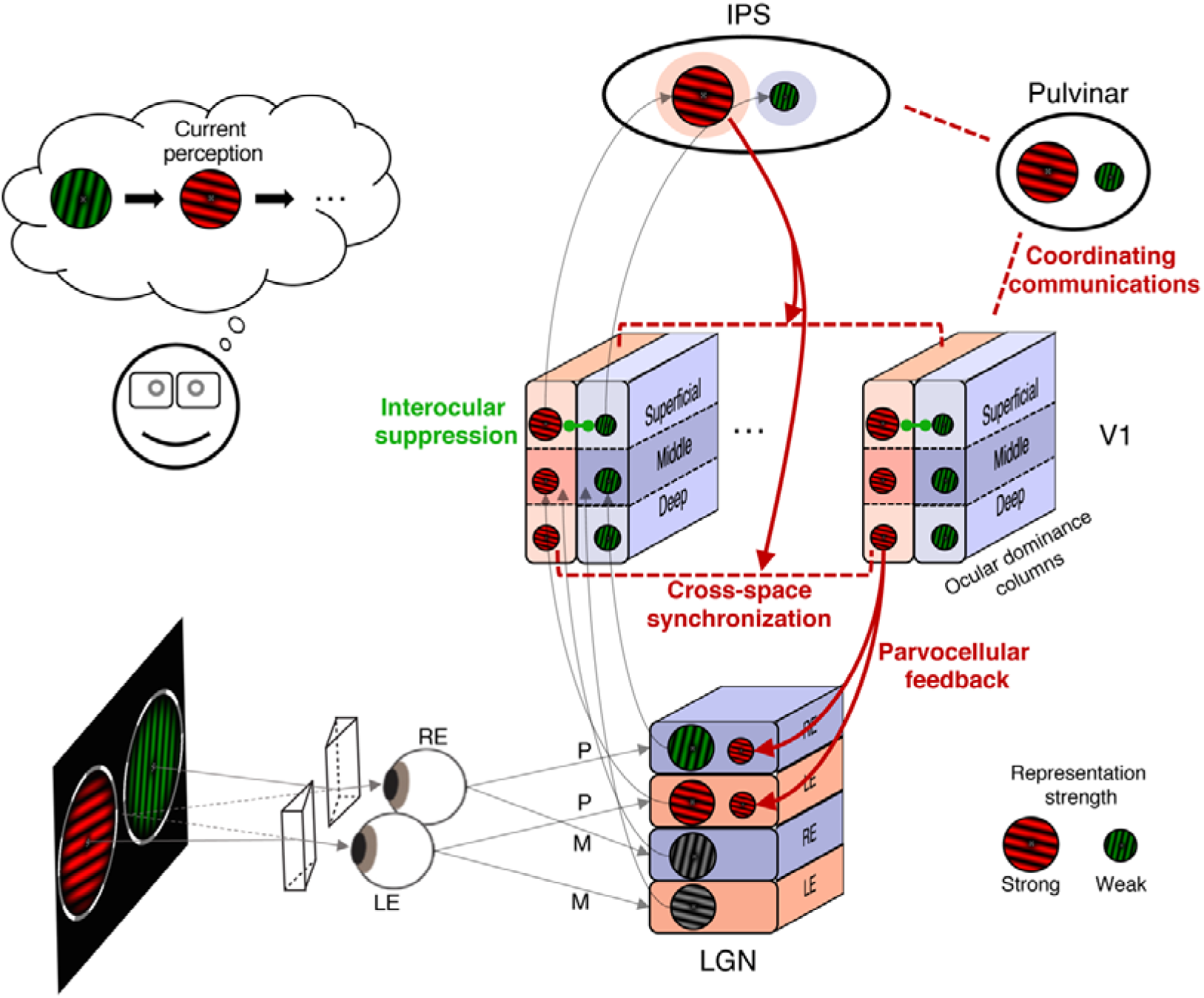

**Highlights:**

- Eye-specific rivalry modulation is strongest in the superficial layers of V1 ODCs and more synchronized in superficial and deep layers
- IPS generates stronger eye-specific response modulation and increases connectivity to V1 during rivalry compared to replay
- LGN activity shows no evidence of eye-specific rivalry modulation but strong perceptual rivalry modulation in its P subdivision
- IPS and ventral pulvinar show robust perceptual rivalry modulation and increased connectivity to the early visual cortex

## Introduction

Two incompatible images presented to the two eyes compete for access to consciousness. This visual illusion, called binocular rivalry, is an ideal model to study how our brain resolves visual ambiguity (R Blake & Logothetis, 2002; Dayan, 1998; Wilson, 2003), a key mechanism to generate conscious visual perception (Randolph Blake et al., 2014; Crick, 1996; Myerson et al., 1981). Although rivalry-related activity has been found in many brain areas (Brascamp et al., 2018; Tong et al., 2006), how hierarchical neural processes in our brain interact to resolve visual competition remains unclear.

It has been proposed that binocular rivalry could arise from interocular competition in early visual areas (R Blake, 1989), either through lateral mutual inhibition between adjacent ocular dominance columns (ODCs) in the primary visual cortex (V1), or interlaminar inhibition between adjacent ocular layers in the lateral geniculate nucleus (LGN) of the thalamus (Kacie Dougherty et al., 2018, 2021; Guillery & Colonnier, 1970). In support of this hypothesis, human fMRI studies found robust eye-specific rivalry modulations in the blindspot area (Tong & Engel, 2001) and ocular-biased voxels in V1 (Haynes et al., 2005), and even in the LGN (Haynes et al., 2005). However, since these early fMRI studies didn’t resolve activity from V1 ODCs or LGN ocular layers, the neural mechanisms of interocular competition still lack concluding evidence. Inconsistent with these fMRI results, single-unit spiking activity showed no evidence of binocular rivalry in the LGN of alert monkeys (Lehky & Maunsell, 1996), and a weak effect in V1 (Leopold & Logothetis, 1996). Since BOLD signals can reflect synaptic input activity (Logothetis & Wandell, 2004), one possible explanation for the discrepancy between single-unit and fMRI results is that feedback modulations from higher-order brain areas drive rivalry-related activity in the early visual areas (de Jong et al., 2020; Maier et al., 2008). A potential role of eye-specific feedback in resolving interocular conflicts is supported by behavioral evidence that top-down attention can be eye-specific (Zhang et al., 2012), and by electrophysiology and neuroimaging evidence of eye-specific representations in extrastriate cortex (Burkhalter & Van Essen, 1986; Maunsell & Van Essen, 1983; Schwarzkopf et al., 2010; Zaretskaya et al., 2020). Therefore, it remains unclear whether interocular competition in binocular rivalry arises from interlaminar inhibition between ocular layers in the LGN, lateral inhibition between V1 ODCs, or is driven by eye-specific feedback from higher-order brain areas.

In addition to interocular competition, binocular rivalry could also involve pattern competition between stimulus representations at multiple levels of the visual hierarchy, and possibly attention and perceptual decision-making mechanisms in high-level brain areas. In this hierarchical whole-brain network, the role of frontoparietal areas is the most debated. Although a causal role of frontoparietal activity in generating perceptual transitions remains controversial (Brascamp et al., 2015; Lumer et al., 1998), converging evidence demonstrate that binocular rivalry requires top-down attention (Brascamp & Blake, 2012; Li et al., 2017; Zhang et al., 2011), suggesting a potential role of the frontoparietal attention network in resolving visual competition. Moreover, whether frontoparietal areas also represent perceptual state during bi-stable perception requires further investigation (Kapoor et al., 2022; Mashour et al., 2020; Tononi et al., 2016). Another subcortical area that is rarely investigated but might play important roles in perceptual rivalry is the pulvinar of the thalamus. Interconnected with frontoparietal areas and visual cortex, pulvinar may regulate information transfer between cortical areas and support cortical computations to resolve perceptual conflicts (Jaramillo et al., 2019; Saalmann et al., 2012; Wilke et al., 2009; Zhou et al., 2016). Finally, parallel visual pathways might be differentially involved in binocular rivalry. Although behavioral studies suggest that the parvocellular (P) pathway is more involved in rivalry than the magnocellular (M) pathway (He et al., 2005), there is no direct neuroimaging evidence supporting this hypothesis. To this date, the hierarchical whole-brain network of perceptual rivalry has not been clearly demonstrated.

Using high-resolution fMRI at 7 Tesla to measure mesoscale activity in the human brain, we investigated hierarchical neural mechanisms underlying binocular rivalry in cortical and subcortical areas. To reveal the neural circuitry of interocular competition, Experiment 1 and 2 studied eye-specific rivalry modulations in V1 ODCs at different cortical depth and LGN ocular layers, and also in the higher-order extrastriate and parietal cortex. To investigate the hierarchical whole-brain network of perceptual rivalry, Experiment 3 used M and P pathway-selective visual stimuli to study perceptual rivalry modulations over the whole brain.

## Results

In the rivalry condition of Experiment 1 and 2, a pair of red/green gratings in orthogonal orientations were dichoptically presented to the two eyes. Subjects reported their perception with button presses (red, green or mixed). In the replay condition, monocular images were presented in alternations with simulated transitions to match the temporal sequence of perception during binocular rivalry, and subjects reported their perception as in the rivalry condition. In localizer runs, a full contrast counterphase flickering checkerboard was alternatively presented to the two eyes to measure the ocular bias of voxels in V1 and LGN.

Based on known anatomical connections of the primate geniculostriate pathway (Felleman & Van Essen, 1991), feedforward input from the LGN mainly terminates in the middle layer (layer 4) of V1, cortico-cortical feedbacks target the superficial (layers 1/2/3) and deep layers (layers 5/6), and lateral inhibition between ODCs through horizontal connections are most prominent in the superficial layers (layers 2/3) (Buzs et al., 2001; K Dougherty et al., 2019; Gilbert & Wiesel, 1983; Sengpiel et al., 1995). If interocular competition arises from interlaminar inhibition in the LGN (fig 1b, first hypothesis), eye-specific rivalry modulation should be strongest in V1 middle layer. Otherwise, if interocular competition arises from lateral inhibition between V1 ODCs (second hypothesis), eye-specific modulation should be strongest in the superficial layers. Finally, if feedback processes are involved to resolve interocular competition (third hypothesis), eye-specific effect of rivalry should be stronger in the superficial and deep layers compared to the middle layer. Although there is no clear evidence that corticogeniculate feedback can be eye-specific, we still included it as a possibility in the last hypothesis according to the (Haynes et al., 2005) study.

**Figure 1.**
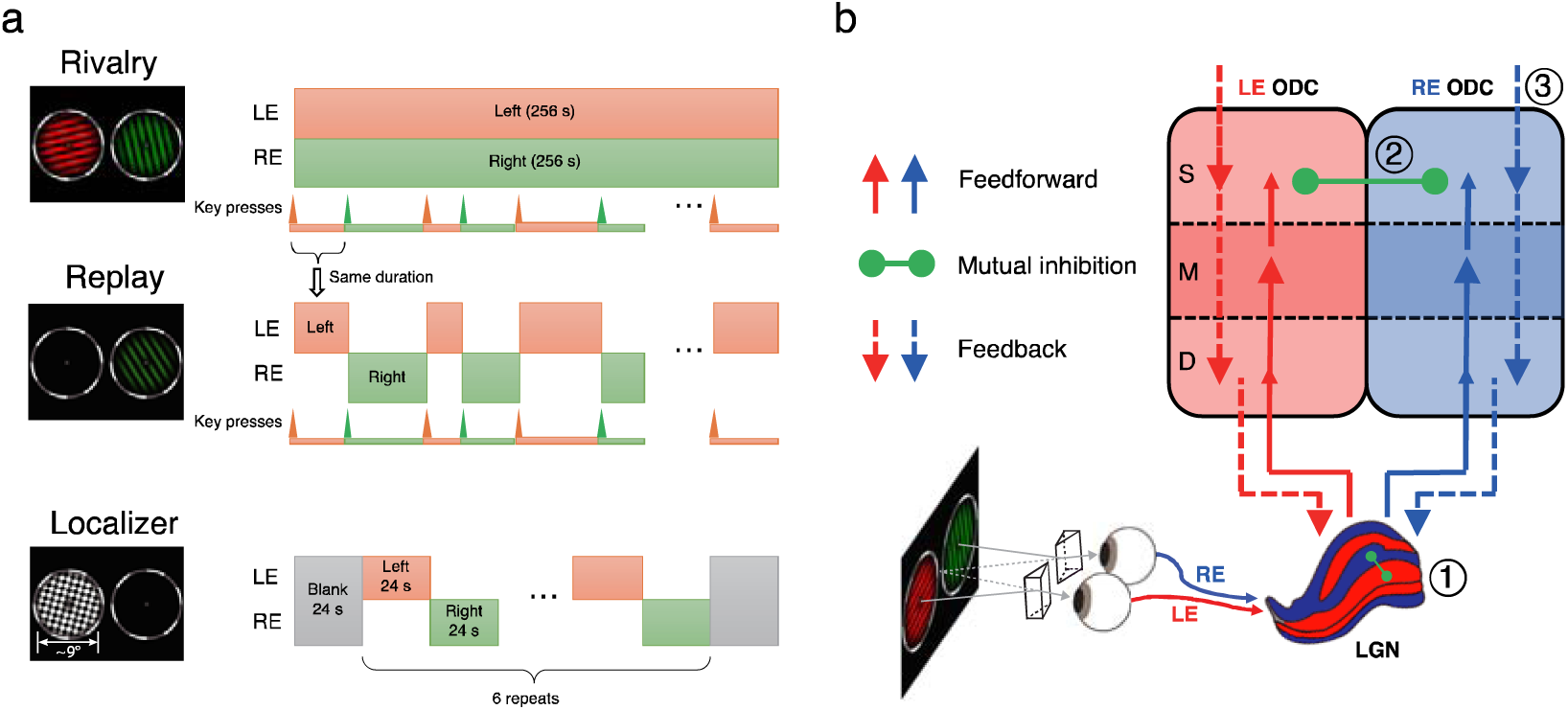
**(a) Stimuli and procedures of Experiment 1 and 2.** In rivalry runs, rotating red and green gratings in orthogonal orientations were dichoptically presented to the two eyes. In replay runs, monocular stimuli were alternatively presented to the two eyes to simulate the perception in the previous rivalry run. In localizer runs, a high contrast flickering checkerboard was monocularly delivered to the two eyes in alternation. **(b) Possible neural circuits of interocular competition in binocular rivalry.** (1) Interlaminar mutual inhibition between LGN ocular layers. (2) Lateral mutual inhibition between V1 ODCs. (3) Eye-specific feedback modulation from higher-order cortical areas. Solid and dashed arrows indicate feedforward and feedback connections, respectively. Green dots connected by solid green lines denote mutual inhibitions. Abbreviations: S (superficial), M (middle), D (deep), LE (left eye), RE (right eye), ODC (ocular dominance column).

### Eye-specific rivalry signal modulation is strongest in the superficial depth of V1 ODCs

In Experiment 1, we tested the three hypotheses using cortical layer-dependent fMRI at submillimeter resolution. T2*w BOLD signals from the early visual cortex and parietal cortex were acquired with a gradient echo planar imaging (GE-EPI) sequence at 0.8-mm isotropic resolution. Interdigitated patterns of V1 ODCs can be robustly resolved (Fig. 2a for a representative subject S01, Fig. S1 for all subjects), consistent with our recent study (de Hollander et al., 2021). The orientations of ODCs are roughly perpendicular to the V1/V2 boundary in its vicinity, and highly reproducible across sessions on different days (Fig. S2, *r* = 0.697, *p* < 0.001, Monte Carlo test). Event-related average of eye-specific modulations were time-locked to button presses reflecting perceptual switches (Fig. 2b). From the time of a perceptual switch, BOLD signals increased when subjects perceived the preferred stimulus of the ocular-biased voxels (LE/RE percept for LE/RE biased voxels), and decreased when the non-preferred stimulus was perceived (RE/LE percept for LE/RE biased voxels). The modulation amplitude during binocular rivalry was about 40% of that during stimulus replay. The map of rivalry modulation (difference between the LE and RE percepts, 8-mm FWHM high-pass filtered) matched well with the ODC map acquired with the localizer (Fig. 2c, *r* = 0.475, *p* < 0.001). These results clearly demonstrate that eye-specific modulation of V1 activity in binocular rivalry occurs at the level of cortical columns.

**Figure 2.**
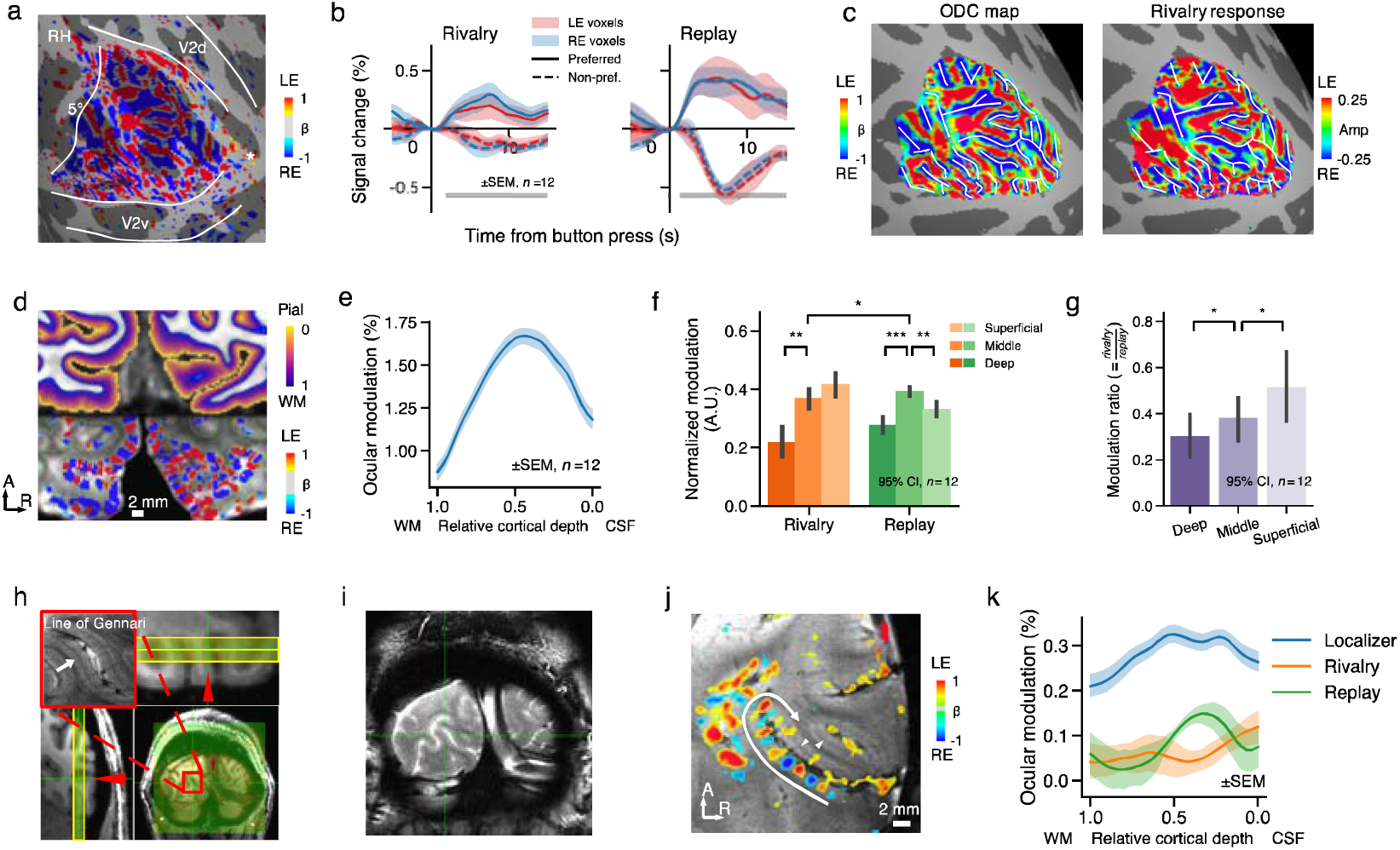
Eye-specific response modulation in V1 ODCs at different cortical depths in Experiment 1. **(a)** OD patterns of a representative subject (S01, the author P.Z.) on the inflated cortical surface of right hemisphere. The color indicates beta values for LE-RE contrast (abs(t) > 2 or *p* < 0.05 uncorrected), same for lower panel in (d). The white lines delineate V1/V2, as well as the 5° eccentricity based on the Benson14 atlas (Benson et al., 2014). **(b)** Event-related timecourses of eye-specific modulation in ocular-biased voxels. Solid (dashed) lines indicate responses to the preferred (non-preferred) percept for the voxel. Error bars indicate SEM across subjects. The bars below indicate time points showing significant differences between different percepts (cluster-based permutation test, cluster defining threshold and cluster-wise FWE corrected p < 0.05) **(c)** Comparison of the ODC map from the ocular-bias localizer and eye-specific response pattern during binocular rivalry for S01. The white reference lines were traced according to the left panel. **(d)** Equivolume cortical depth map overlaid on the T1-weighted image for S01 (upper); GLM beta map (LE-RE) within the gray matter overlaid on the mean EPI image (lower). Purple and green lines indicate the pial and white matter surfaces, respectively. **(e)** Eye-specific response modulation peaked at intermediate depth in the ocular-bias localizer. **(f)** Normalized eye-specific modulation from different cortical depths in rivalry and replay conditions. Error bars indicate 95% confidence interval from bootstrap. **(g)** The modulation ratio of rivalry and replay conditions. **(h)** Slice prescriptions for the 2D-bSSFP experiment (0.5 mm in-plane resolution, 3 mm thickness, perpendicular to the surface) in a representative subject (S06, the author C.Q.). From the T2*-weighted GRE image (upper left inset), the line of Gennari is clearly visible in the middle layer of V1 gray matter. **(i)** A raw bSSFP image frame. **(j)** ODCs can be clearly identified on the cross section of calcarine sulcus (white arrow). **(k)** The V1 depth profile of eye-specific modulation in the rivalry, replay and localizer conditions from T2-weighted BOLD signals with bSSFP fMRI. Shaded areas indicate SEM across runs (13 runs for the localizer, and 6 runs each for the rivalry and replay conditions).

Cortical depth was estimated for each voxel with an equivolume method (Waehnert et al., 2014), based on manually edited cortical surface reconstructions (Fig. 2d upper). The columnar structure of ODCs perpendicular to the cortical surface can be clearly seen (Fig. 2d lower). The differential response between the left and right eye stimulation in the localizer peaked in the middle depth of V1 (Fig. 2e), consistent with the fact that thalamocortical projections terminate mainly at layer 4C. Thus, the OD column-specific response derived from the differential of balanced responses to the LE and RE stimuli largely reduced the non-specific signals in the superficial layers associated with the blooming effect of pial veins (Moon et al., 2007; Uludag & Havlicek, 2021). We next investigated how this laminar-columnar circuit reflects the endogenous mechanisms that gate perceptual awareness in binocular rivalry. Normalized eye-specific modulations in the rivalry and replay conditions are shown in Fig. 2f. Two-way repeated measures ANOVA showed a significant interaction of cortical depth (superficial/middle/deep) and stimulus conditions (rivalry/replay): *F*(2,22) = 6.372, *p* = 0.013, 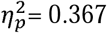. In the replay condition, the middle layer showed the strongest effect of eye-specific modulation (main effect of depth, *F*(2,22) = 15.290, *p* < 0.001 Bonferroni corrected, 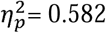; deep vs. middle, *t*(11) = -6.218, *p* < 0.001, Cohen’s *d* = 2.921; middle vs. superficial, *t*(11) = 3.608, *p* = 0.004, Cohen’s *d* = 1.600), consistent with the feedforward input from the LGN. During binocular rivalry, eye-specific modulation was more biased to the superficial depth (main effect of depth, *F*(2,22) = 12.859, *p* < 0.001, 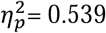; deep vs. middle, *t*(11) = -3.486, *p* = 0.005, Cohen’s *d* = 1.794; middle vs. superficial, *t*(11) = -1.617, *p* = 0.134, Cohen’s *d* = 0.665). The laminar profiles without normalization were qualitatively the same. To directly reveal the difference in depth profile between the two conditions, we calculated a modulation ratio by dividing the rivalry modulation by the replay modulation. This rivalry/replay modulation ratio was strongest in the superficial layer (Fig. 1g; main effect of depth, *F*(2,22) = 8.118, *p* = 0.009, 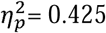; deep vs. middle, *t*(11) = -2.403, *p* = 0.035, Cohen’s *d* = 0.479; middle vs. superficial, *t*(11) = -2.551, *p* = 0.027, Cohen’s *d* = 0.578).

One subject also performed multiple sessions of the same experiment using a passband bSSFP sequence. Compared to the T2*w GE-BOLD signals, T2w BOLD signals from bSSFP fMRI are more sensitive to microvasculature activity in the gray matter, which is closer to the site of neural activity. Previous studies show that T2w BOLD has higher spatial specificity to reveal the laminar profile of cortical processing (Beckett et al., 2020; Liu et al., 2020; Olman et al., 2012; Scheffler et al., 2018). Two coronal slices (0.5-mm in-plane resolution with 3-mm slice thickness) were carefully prescribed to be perpendicular to the calcarine sulcus in one hemisphere, where the ODCs went approximately parallel with the orientation of ‘pencil’ voxels (Fig. 2h). One of the slices shows clear ODC patterns (Fig. 2j), confined within gray matter and highly reproducible across sessions (Fig. S2). Eye-specific modulation peaked in the middle layer in the replay and localizer conditions, but in the superficial layer during rivalry (Fig. 2k). This laminar pattern is consistent with the GE-EPI data. This finding further supports the notion that interocular competition in binocular rivalry mainly arises from lateral mutual inhibitions between ODCs in V1 superficial layers.

### Eye-specific rivalry dynamics is more synchronized in V1 superficial and deep layers

Local interocular competition may result in different local winners and piecemeal perception. It has been hypothesized that feedback signals from higher order areas help synchronize and stabilize local competitions into a globally coherent perceptual state over extended visual field (Kovács et al., 1996; Tong et al., 2006). To test this hypothesis, we characterized the synchrony of eye-specific modulations across V1 ODCs by calculating TR-by-TR Pearson correlations between the ongoing V1 response pattern and the localizer-derived OD pattern (Fig. 3a). More synchronized OD dynamics would predict larger correlation coefficients, quantified by the width of their distribution (O’Hashi et al., 2018; Omer et al., 2018). As stimuli-driven responses were fully coherent in the replay condition, this gives us a benchmark against which to compare the rivalry response patterns at each cortical depth.

**Figure 3.**
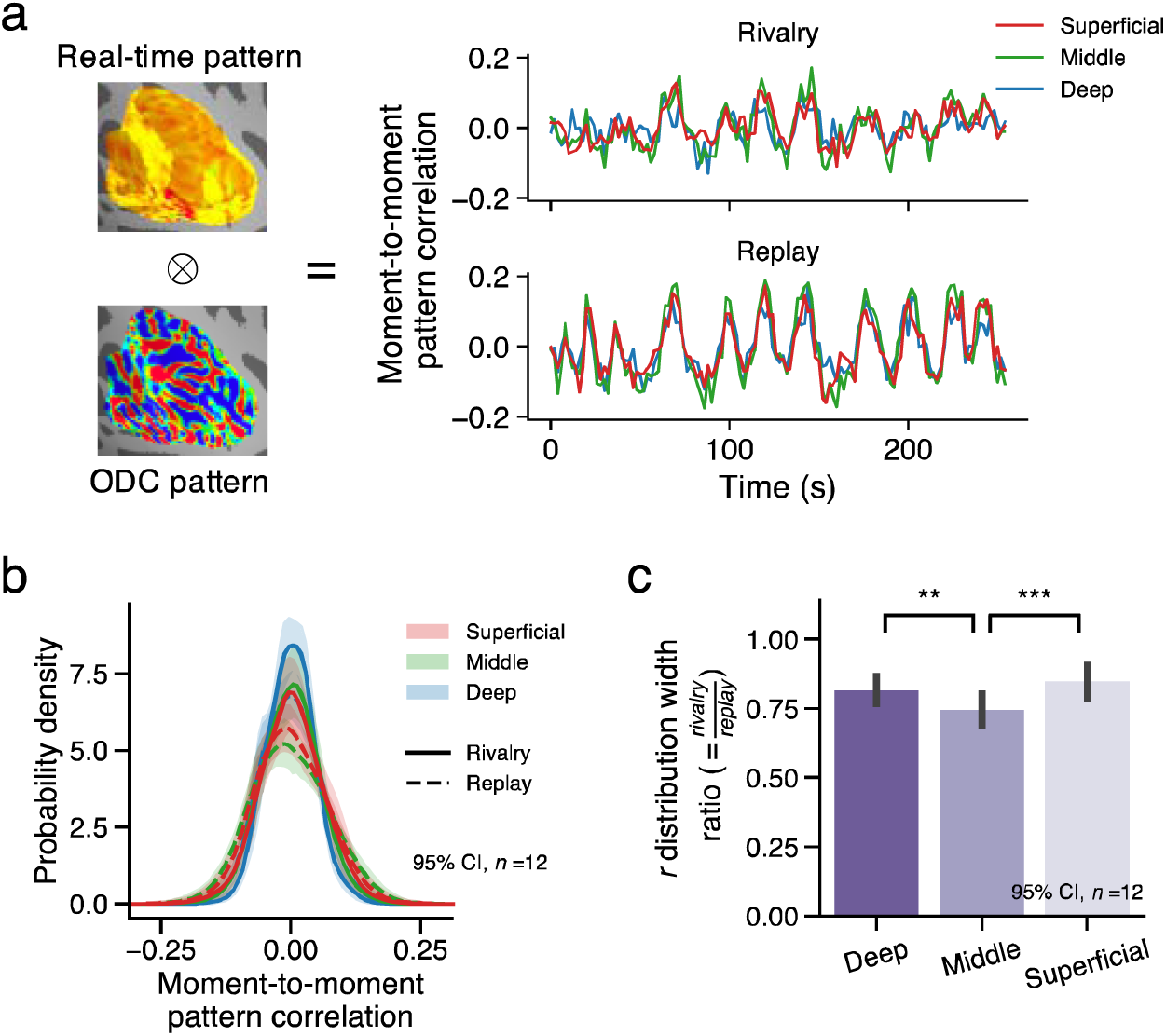
Pattern synchronization across ODCs in different cortical depth. **(a)** TR-by-TR Pearson correlations between the OD pattern from the localizer and the real-time V1 response pattern in typical runs of rivalry and replay. **(b)** The distributions of pattern correlation (r) for different V1 layers in rivalry (solid) and replay (dashed). Shaded area indicates 95% confidence intervals across subjects. **(c)** The ratio of r distribution widths between rivalry and replay conditions across V1 layers.

As predicted, we found that the replay condition was associated with a larger distribution width of correlation coefficients (Fig. 3b). The critical question is, how does the synchrony of OD dynamics differ across cortical depth during rivalry? If pattern synchronization relies on feedback signals, its signature would be more evident in the superficial and deep layers. Indeed, V1 superficial and deep layers showed significantly larger normalized distribution width than the middle layer (Fig. 3c; main effect of depth, *F*(2,22) = 11.789, *p* < 0.001, 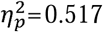; deep vs. middle, *t*(11) = 3.210, *p* = 0.008, Cohen’s *d* = 0.690; middle vs. superficial, *t*(11) = -4.789, *p* < 0.001, Cohen’s *d* = 0.942). To verify whether this actually reflected a difference in the temporal structure of the response or a mere SNR difference across cortical depths, we performed a permutation analysis. A GLM with variable duration of perceptual states was fitted to the vertex timeseries in the rivalry and replay conditions, and the residuals were temporally permuted independently for each vertex before being recombined with the fitted timeseries. The permutation destroyed any synchronous fluctuation unmodeled in the GLM without changing the overall SNR across layers, which we reasoned might reduce the observed laminar difference. As expected, the difference between the deep and middle layers was largely eliminated for the permuted data (Fig. S3; *t*(11) = 0.389, *p* = 0.705, BF_01_ = 3.257, Cohen’s *d* = 0.038) and the main effect of depth was no longer significant (*F*(2,22) = 2.115, *p* = 0.165, 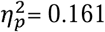). These results suggest that feedback processes may be involved to synchronize local interocular competitions in V1 into a spatially coherent visual representation. We next turned to a prime candidate for the source of these feedback signals, the IPS region of the parietal lobe.

### IPS generates stronger eye-specific response modulation and increased connectivity to the early visual cortex during rivalry compared to replay

As a candidate source of the feedback signals, the intraparietal sulcus (IPS) region of the attention network was suggested to play critical roles in bi-stable perception by TMS studies (Carmel et al., 2010; Kanai et al., 2011; Zaretskaya et al., 2010). Moreover, attention is necessary for binocular rivalry (Brascamp & Blake, 2012; Zhang et al., 2011), and top-down attention can be eye-specific (Zhang et al., 2012). Therefore, we first examined whether IPS encoded the state of currently dominating eye, and its relationship with rivalry modulations in V1. Exploiting the sensitivity of multivariate methods, we trained a support vector machine (SVM) to predict the eye-of-origin of stimulus using data from the ocular-bias localizer. Voxels were selected based on their visual responsiveness and ocular bias (see methods). The model was then used to predict the eye-of-origin of the perceived stimulus in a TR-by-TR basis during rivalry and replay. The distance between each activation pattern and the SVM decision boundary was used as a graded measure of the eye-of-origin representation (Fig. 4a/b). Event-related averages around the time of perceptual switches showed that, in IPS, significant eye-specific modulation was observed in binocular rivalry but not in stimulus replay (Fig. 4c/d; cluster-based permutation test), similar in its posterior (pIPS, IPS0-2 in (Wang et al., 2015)) and anterior (aIPS, IPS3-5) portions (condition*ROI interaction not significant, *F*(1,11) = 0.238, *p* = 0.635, 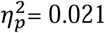). It suggests that IPS might play a role in resolving interocular conflicts, e.g. by setting a competition bias to one eye. Indeed, the modulation amplitudes of IPS significantly correlate with those of V1 only in the rivalry condition (Fig. 4e; rivalry *r* = 0.778, *p* = 0.003; replay *r* = 0.409, *p* = 0.186; and their difference marginally significant, *p* = 0.055 assessed using bootstrap). Note that we tried to avoid the mere influence of inter-subject variation in switch duration by taking into account the duration of perceptual states in the GLM when estimating the modulation amplitude.

**Figure 4.**
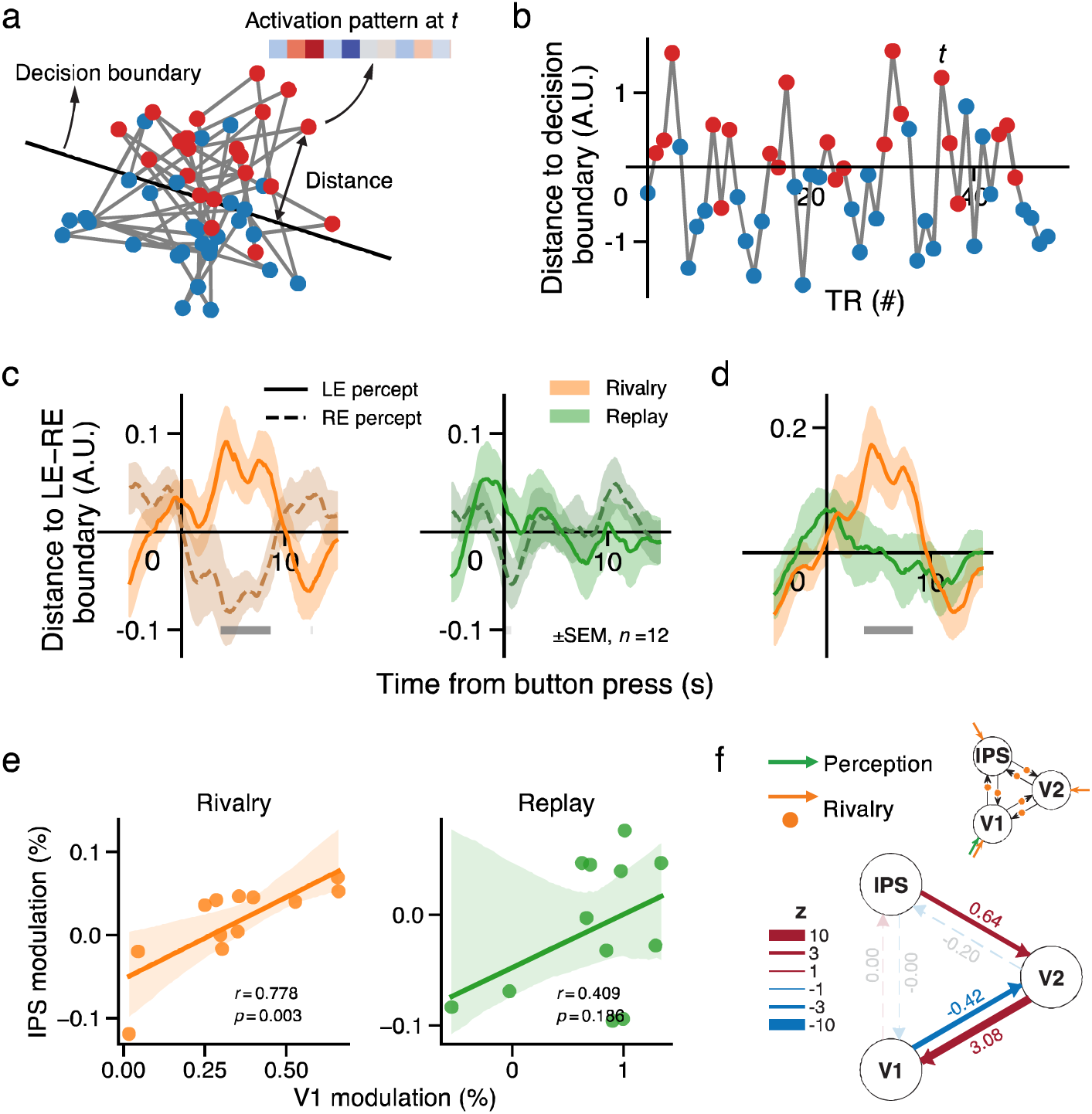
Eye-specific modulations in IPS and its relationship with the early visual cortex. **(a)** Schematic about the distance of activation pattern at each time point to the SVM decision boundary for LE vs. RE stimulation. Red (blue) dots denote the activation patterns of TRs when LE (RE) was stimulated. **(b)** The same distances replotted as a timecourse, i.e., a multivariate differential response. **(c)** Event-related average of the decoding-distance timecourse in IPS for LE (solid) and RE (dashed) percepts in rivalry (orange) and replay (green). **(d)** Differential waveforms between LE and RE events. The gray bars indicate a significant difference between rivalry and replay. Shaded area indicates SEM. Light and dark gray bars in (c) and (d) denote time points with significant difference from zero for corresponding conditions before and after multiple testing correction, respectively. **(e)** Inter-subject correlation between eye-specific modulations in IPS and V1. Shaded area indicates 95% confidence interval of the linear fit. **(f)** Changes in effective connectivity among V1, V2, and IPS during rivalry compared to replay. Numbers beside the connections denote the estimated modulatory effects in coupling strength, and the line thickness indicate the z value. Non-significant connections (*P_p_* < 0.95) were rendered as faint, dashed lines. Inset: DCM model specification for effective connectivity between IPS, V2 and V1. Green arrow: Eye-specific driving input for both rivalry and replay. Orange arrows: additional eye-specific driving inputs for rivalry only. Orange dots: eye-specific modulatory effects of rivalry.

Eye-specific rivalry modulation was also found in V2 of the extrastriate cortex (Fig. S4). To further investigate the causal relationship of eye-specific activity between V1, V2, and IPS, we performed a dynamic causal modeling (DCM) analysis on the multivariate projected timeseries that best discriminated LE from RE perception based on SVM trained on localizer runs. In the full DCM model (Fig. 4f, upper panels), V1 receives eye-specific driving inputs in both rivalry and replay conditions. Intrinsic or fixed connections were defined between and within cortical areas. The between-area connections as well as each node could be modulated or driven by an additional eye-specific input in rivalry but not in replay. This input thus captured the difference between the two conditions. The full DCM model was estimated for each individual (Zeidman, Jafarian, Corbin, et al., 2019). For group-level analysis, we used Parametric Empirical Bayes (PEB), Bayesian model reduction, and Bayesian model average to make inference about the model parameters (Friston et al., 2016; Zeidman, Jafarian, Seghier, et al., 2019). Compared to stimulus replay, binocular rivalry significantly increased the feedback connectivity from IPS to V2 and from V2 to V1 (posterior probability (*P_p_*) = 0.997 and 1.000, respectively), whereas decreased the feedforward connectivity from V1 to V2 (*P_p_* = 0.999) and the driving input to V1 (*P_p_* = 1.000). These findings support IPS as the source of feedback processes in resolving and synchronizing interocular competitions in V1 ODCs.

### No evidence for eye-specific rivalry modulation in ocular-biased clusters of the LGN

In Experiment 2, to investigate whether the LGN was involved in interocular competition during binocular rivalry, we used a 1.2-mm isotropic GE-EPI sequence to study ocular-layer selective signals in the LGN. Stimuli and procedures were similar as in Experiment 1, fMRI slices were orientated to cover both LGN and V1. Robust ocular-biased patterns were clearly and consistently revealed in the LGN across sessions in separate days by the ocular-bias localizer (Fig. 5a/b for S01; *r* = 0.862, *p* < 0.001, Monte Carlo test, see Fig. S5 for the ocular-biased pattern of all participants). There are two ocular-biased clusters for each LGN: a ventromedial one biased to the ipsilateral eye, and a dorsolateral one biased to the contralateral eye, which replicated the findings of our recent study (Qian et al., 2020). Based on the simulation analysis of the previous study, these ocular-biased clusters were results of BOLD blurring and fMRI down-sampling of the LGN laminar pattern (shown here in the lower right of Fig. 5a). Therefore, BOLD signals from the ocular-biased clusters represent ocular layer-selective activity of the LGN. For more details about the simulation analysis, please refer to figure 1b and figure 3a in our previous study (Qian et al., 2020).

**Figure 5.**
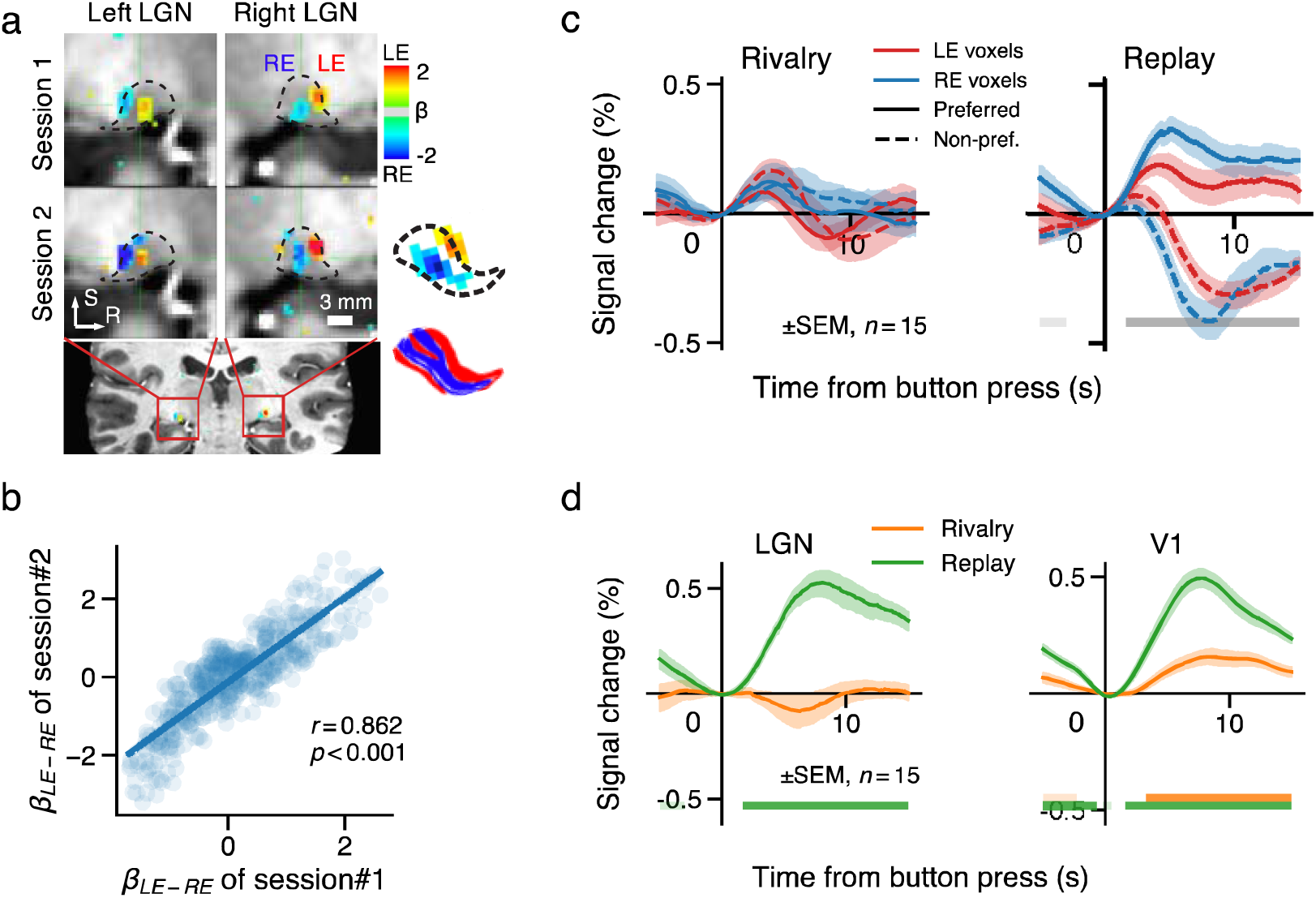
Eye-specific modulation in ocular-biased clusters of the LGN. **(a, b)** Highly reproducible ocular-biased clusters in the LGN (LE-RE beta maps, p < 0.01 uncorrected) of a representative subject (S01). See Fig. S5 for all participants. Inset: simulation results of ocular-biased clusters, reproduced from Fig. 3a in (Qian et al., 2020). **(c)** Event-related average timecourses for preferred and non-preferred percepts in the LGN ocular-biased clusters. **(d)** Eye-specific modulation timecourses in LGN and V1 during rivalry and replay. Horizontal bars in light and dark color denote time points with significant difference from zero for corresponding conditions before and after multiple testing correction, respectively.

To our surprise, although BOLD signals in ocular-biased clusters of the LGN showed a transient increase after perceptual switches (Fig. 5c, mean response to both preferred and non-preferred switches averaged between 2-6 s, *t*(14) = 4.112, *p* = 0.001, Cohen’s *d* = 1.062), there was no significant eye-specific modulation during binocular rivalry (Fig. 5d; *t*(14) = -0.619, *p* = 0.546, BF_01_ = 3.226). As positive controls, the differential response to preferred- and non-preferred eye stimulation averaged between 4-12 s was as strong in LGN as in V1 in the replay condition, and V1 showed robust eye-specific modulations in both conditions (Fig. 5d). Although there was no evidence of eye-specific rivalry modulation in the ocular-biased clusters, a negative eye-specific effect can be found in LGN voxels outside the ocular-biased clusters (Fig. S6). This negative effect is likely due to attentional suppression outside the stimulus region (Shmuel et al., 2002; Tootell et al., 1998). No significant eye-specific modulation was found in the pulvinar (*t*(14) = 1.504, *p* = 0.155, Cohen’s *d* = 0.388).

Therefore, our 7T fMRI results suggest weak, if any, eye-specific rivalry modulation within the stimulus regions of the LGN. This finding is consistent with the single-unit study of macaque LGN (Lehky & Maunsell, 1996), and also with the observation in Experiment 1 that eye-specific rivalry modulation peaked in V1 superficial layers. If interocular competition of binocular rivalry was resolved in the LGN, one would expect a peak of eye-specific modulation in the middle layer of V1 that receives thalamic input. Although it is unlikely that feedback processes can modulate ocular-layer selective activity of the LGN in binocular rivalry, it remains possible that perception-related feedback processes can modulate LGN activity in a non-eye-specific manner (Wunderlich et al., 2005). To test this hypothesis and to investigate the whole-brain network of stimulus-specific or perceptual rivalry modulation, we conducted a third experiment.

### Robust perceptual rivalry modulation in P subdivisions of the LGN and ventral pulvinar

In Experiment 3, we exploited the spatial segregation of magnocellular (M) and parvocellular (P) pathways in subcortical (e.g., LGN, ventral pulvinar) and cortical (e.g, MT+, hV4) visual areas as a novel tagging method to study the effect of perceptual rivalry across the whole brain. According to the laminar organization of the LGN, M and P layers are located in its ventral and dorsal portions, respectively. In the ventral pulvinar (vPul), the parvocellular lateral portion reciprocally connects with the early visual cortex and ventral visual stream, while the magnocellular medial portion receives input from the superior colliculus (SC) and connects with dorsal visual stream such as area MT (Arcaro et al., 2015; Bridge et al., 2016; Kaas & Lyon, 2007).

We designed M- and P-biased visual stimuli to preferentially activate the M and P visual pathways (Derrington & Lennie, 1984; Wiesel & Hubel, 1966). The M stimulus was a low spatial frequency (0.5 c.p.d.) achromatic grating (30% or 50% contrast), while the P stimulus was a red/green equiluminant chromatic grating presented at high contrast (Fig. 6a). M and P gratings were dichoptically presented in orthogonal orientations to the two eyes in binocular rivalry, and monocularly presented to the two eyes in alternation in simulated replay. An independent localizer was used to measure the M-P bias of voxels. We defined the ROIs of M- and P-biased voxels in the LGN and vPul based on their M-P contrast in the functional localizer and anatomical locations (Fig. 6b, see methods for details). The M and P stimuli also selectively activated the dorsal and ventral cortical streams (Fig. S9a).

**Figure 6.**
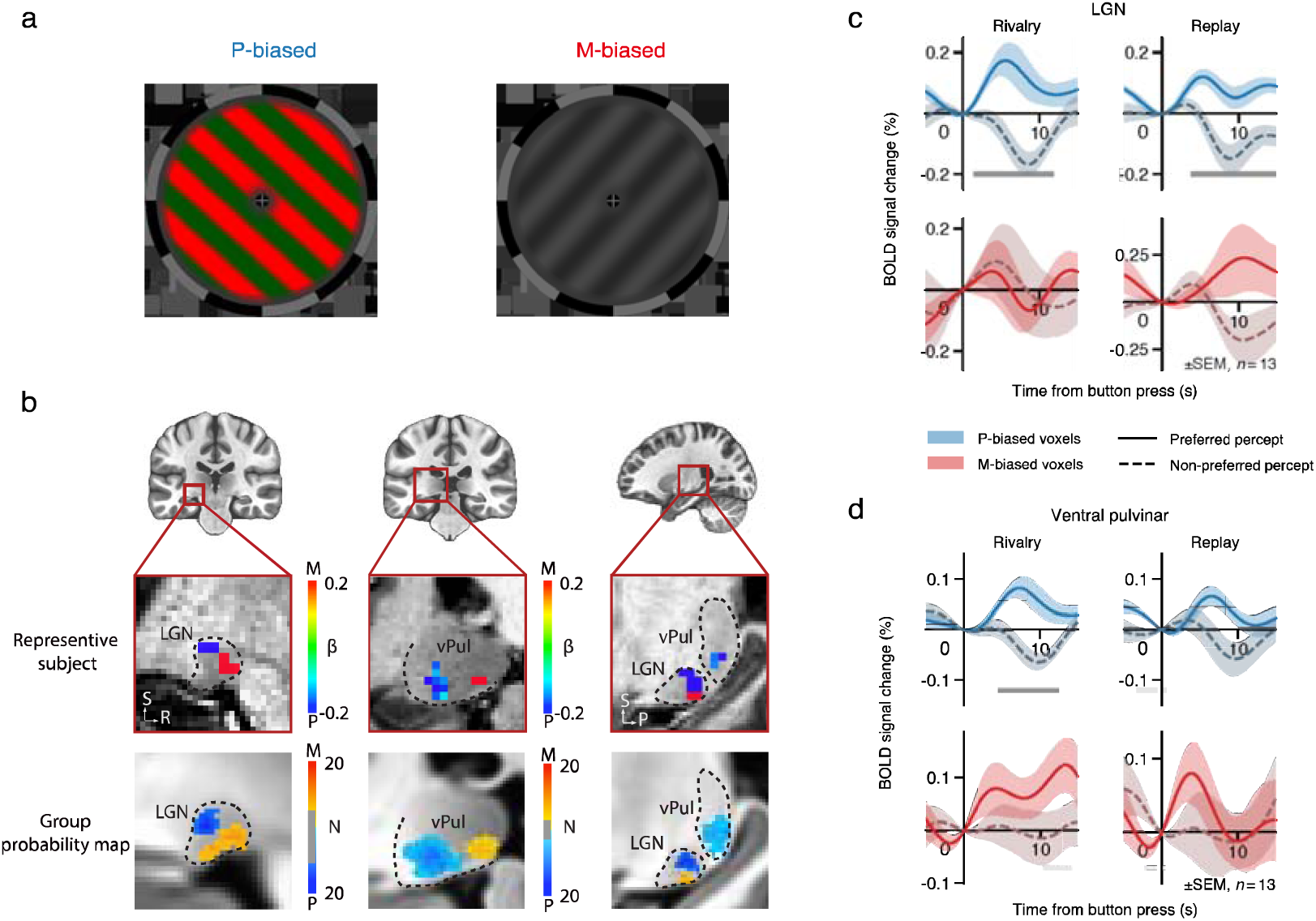
Effect of perceptual rivalry between chromatic and achromatic gratings in the LGN and ventral pulvinar. **(a)** The M-biased achromatic grating and P-biased chromatic grating in Experiment 3. **(b)** Upper: Localizer M-P beta values in M- and P-biased ROIs in the LGN and vPul of a representative subject (S02). The ROIs were defined based on both their M-P response bias in the functional localizer and anatomical locations. Lower: Overlap probability map for M and P ROIs in MNI space. Colors indicate the number of overlaps across subjects and hemispheres (thresholds: LGNp=11, LGNm=5, Pulp=3, Pulm=3). Dashed lines indicate the anatomical boundaries of LGN and pulvinar. **(c, d)** Event-related BOLD responses in M- and P-biased voxels of the LGN and vPul. Shaded areas represent SEM across subjects. Light and dark gray bars denote time points with significant difference between the two percepts before and after multiple testing correction, respectively.

During binocular rivalry, BOLD signals in P-biased voxels of the LGN increased when subjects reported seeing the chromatic grating, and decreased when the achromatic grating took dominance (Fig. 6c; Fig. 7a, *t*(12) = 2.993, *p* = 0.011 Holm corrected, Cohen’s *d* = 0.830). Similar modulation was also observed in the replay condition (Fig. 7a, *t*(12) = 3.892, *p* = 0.004, Cohen’s *d* = 1.079). For the M-biased voxels, there were marginally significant stimulus-driven modulations in the replay condition (Fig. 7b, *t*(12) = 1.460, *p* = 0.085 uncorrected, Cohen’s *d* = 0.405), but without significant perceptual modulation during binocular rivalry (Fig. 6c, Fig. 7b). The different involvement of LGNp and LGNm in perceptual rivalry were further supported by a significant interaction between condition and pathway (*F*(1,12) = 5.339, *p* = 0.039, 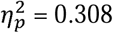 = 0.308). Interestingly, the rivalry/replay modulation ratio in LGNp was significantly larger than V1p (condition*ROI interaction, *F*(1,12) = 7.518, *p* = 0.018, 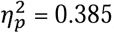), whereas comparable with hV4 (interaction not significant, *F*(1,12) = 0.365, *p* = 0.557, 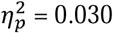), suggesting that the perceptual rivalry modulation in LGNp might be related to feedback signals from higher-order brain areas.

**Figure 7.**
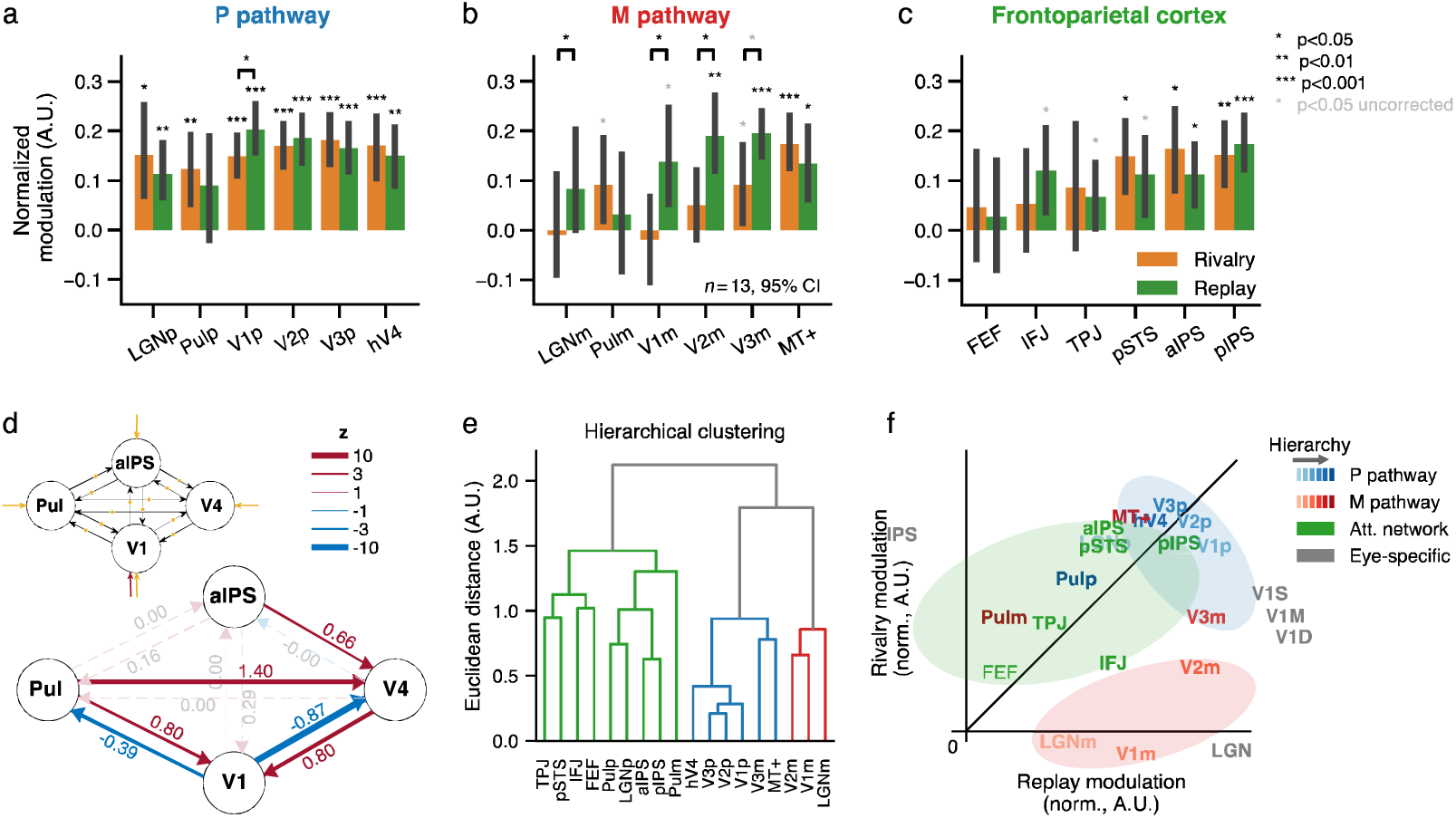
Hierarchical whole-brain network for perceptual rivalry. **(a-c)** Normalized rivalry and replay modulations in the P and M visual pathways and frontoparietal areas. The data vector comprising modulations from all subjects and both conditions were normalized to have length one. The p-values were Holm corrected for multiple comparisons. **(d)** Effective connectivity between aIPS, V4, V1 and ventral pulvinar. **(e)** Hierarchical clustering of normalized rivalry and replay modulations in cortical and subcortical areas. **(f)** 2D visualization of mean-normalized rivalry and replay modulations of all ROIs.

For the ventral pulvinar, both P- and M-biased voxels showed significant perceptual response modulations during binocular rivalry (Fig. 6d; Fig.7a for Pulp, *t*(12) = 3.347, *p* = 0.009, Cohen’s *d* = 0.928; Fig. 7b for Pulm, *t*(12) = 2.079, *p* = 0.030 uncorrected, Cohen’s *d* = 0.576), whereas only P-biased voxels showed marginally significant stimulus-driven modulations in simulated replay (*t*(12) = 1.596, *p* = 0.068 uncorrected, Cohen’s *d* = 0.443). Unlike the LGN, no significant interaction was found between condition and pathway (*F*(1,12) = 0.072, *p* = 0.793, 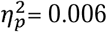). Searchlight decoding also revealed stimulus-related representations in the ventral pulvinar in both rivalry and replay conditions (Fig. S9c). We did not find significant perceptual state-related modulation in the SC in either rivalry or replay (Fig. S8c), thus SC data were not included in the following analysis.

### Perceptual rivalry modulation is weak or absent in early visual areas of the magnocellular pathway

Robust perceptual rivalry modulation can be found in P-biased voxels of the early visual cortex (V1p, V2p, V3p, hV4, Fig. 7a). Perceptual modulation in binocular rivalry was significantly smaller than stimulus-evoked modulation in simulated replay in V1p (t(12) = -3.389, *p* = 0.032, Holm corrected across ROIs, Cohen’s d = 0.602), and the ratio between the response amplitudes of the two conditions steadily went up along the cortical hierarchy (slope = 0.137, 95% CI = [0.051,0.279], *p* = 0.001), consistent with previous reports (Leopold & Logothetis, 1996; Mo et al., 2022). In contrast, early stages of the M pathway did not exhibit significant rivalry modulation (for LGNm, *t*(12) = -0.195, *p* = 0.576, Cohen’s *d* = 0.054; for V1m, *t*(12) = -0.437, *p* = 0.665, Cohen’s *d* = 0.121), but showed significantly stronger modulations in replay than in rivalry (LGNm: *t*(12) = -3.180, *p* = 0.040, Cohen’s *d* = 0.478; V1m: *t*(12) = -2.936, *p* = 0.049, Cohen’s *d* = 0.896). The rivalry/replay modulation ratio also increased along the visual hierarchy (V1m, V2m, V3m, MT+, Fig. 7b), but more than 3-fold faster than the P pathway (*slope* = 0.450, 95% CI = [0.196,1.051], *p* = 0.001). Rivalry and replay modulations were comparable at high-level areas of the M and P pathways (i.e., area MT+ and hV4). Therefore, perceptual rivalry modulation was strong in the P pathway, but weak or absent in early stages of the M pathway.

### Robust perceptual rivalry modulations in IPS and pSTS

We further investigated whether perception-state related information was represented in the frontal and/or parietal association cortices. Since there is no known large-scale segregation of M/P representations in frontoparietal areas, we again deployed multivariate methods to test if perceptual content can be decoded TR-by-TR from these high-order brain areas. Regions with significant visual response in the localizer runs were included (FEF, IFJ, TPJ, pSTS, aIPS, pIPS, see Fig. S7 for ROI definition). To improve decoding performance in these high-level brain areas, a cross-validated grid-search approach was used for feature selection based on visual responsiveness and M/P bias in localizer runs (see methods). The anterior and posterior portions of IPS were engaged differently in rivalry and replay (Fig. 7c, significant ROI *condition interaction in normalized modulation, *F*(1,12) = 8.385, *p* = 0.013, 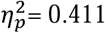). Modulation amplitudes were comparable between rivalry and replay in pIPS, whereas marginally larger in rivalry compared to replay in aIPS (*t*(12) = 2.057, *p* = 0.062 uncorrected, Cohen’s *d* = 0.367; most significant around 8 s after perceptual switch, see Fig. S8b). Similar results were also found in pSTS, showing significant modulation in rivalry (*t*(12) = 3.680, *p* = 0.016, Cohen’s *d* = 1.021), but not in replay after correction (*t*(12) = 2.584, *p* = 0.076, Cohen’s *d* = 0.717). These findings are consistent with the stronger eye-specific rivalry modulation in IPS found in Experiment 1 (Fig. 4). No reliable above-chance decoding was observed in the frontal lobe. Searchlight decoding also revealed significant activations in the parietal lobe during binocular rivalry (Fig. S9b). In sum, perceptual states in binocular rivalry can be decoded from parietal but not frontal activity in our data.

### IPS and ventral pulvinar show stronger perception-state related connectivity to the early visual cortex in rivalry compared to replay

To investigate the information flow of perception-related signals in a whole-brain network, we performed an effective connectivity analysis using DCM within a minimal network containing early and high-level visual cortices of the P pathway (V1, V4), and the parietal cortex (aIPS) and ventral pulvinar (Pul). Here we selected aIPS as the potential source of feedback signals because it showed stronger modulation in rivalry compared to replay. Similar to the eye-specific connectivity in Experiment 1, binocular rivalry showed an enhanced feedback connectivity from aIPS to V4 (*P_p_* = 0.966) and from V4 to V1 (*P_p_* = 1.000) compared to stimulus replay. In addition, connections from Pul to V1 and V4 were also significantly stronger during rivalry compared to replay (*P_p_* = 0.995 and 1.000, respectively). Feedforward connections were significantly weaker during rivalry from V1 to V4 and Pul (*P_p_* = 1.000 for both V1-V4 and V1-Pul connections). These results suggest aIPS as the potential source of feedback signals to the early visual cortex to help resolve perceptual conflicts, e.g., by setting a bias in the competition. Meanwhile, pulvinar may regulate the feedback connectivity across the hierarchy.

### Hierarchical whole-brain network of perceptual rivalry

Finally, to further illustrate the relationship of cortical and subcortical responses, we performed a clustering analysis of rivalry and replay modulations across different brain areas. Each ROI was represented as a vector comprising normalized modulation amplitude during rivalry and replay for each subject, which captured similarities in both rivalry/relay modulation ratio and inter-subject response correlation . Hierarchical clustering procedure based on Euclidean distance suggested the ROIs were best described as 3 groups (Fig. 7e), which was visualized by plotting the ROIs in a 2D-plane according to their normalized modulation amplitude in rivalry and replay (Fig. 7f), along with eye-specific modulation of LGN, V1, and IPS in Experiment 1 and 2. The first group, comprising mainly early stages of the M pathway (LGNm, V1m, V2m), was characterized by the near absence of rivalry modulation and robust replay modulation. Thus, low-level areas of the M pathway are stimulus-driven but modulated little by perceptual rivalry. The second group, including cortical areas of the P pathway and higher levels of the M pathway (V1p, V2p, V3p, hV4, V3m, MT+), featured more or less similar modulation in both conditions. Perceptual modulation in this group became increasingly indistinguishable with stimulus-driven response along the visual hierarchy. The third group mainly composed of cortical and subcortical attention network (aIPS, pIPS, pSTS, TPJ, FEF, IFJ, Pulp, Pulm). Overall, they showed a trend of stronger response modulation in rivalry compared to replay, suggesting a role in resolving competition between conflicting representations. Interestingly, the earliest stage of the P pathway (LGNp) also belonged to this group, consistent with a feedback modulation from higher order cortical areas. These analyses summarize the hierarchical whole-brain network of perceptual rivalry, which may suggest distinct roles of sub-networks in visuosensory processing, perceptual representation and conflict resolution.

## Discussion

Here we studied hierarchical neural mechanisms of binocular rivalry in human cortical and subcortical areas using high-resolution fMRI at 7 T. Experiment 1 and 2 investigated the neural circuitry for interocular competition underlying binocular rivalry. Results show that eye-specific rivalry modulation was strongest in the superficial layers of V1 ODCs, and more synchronized in the superficial and deep layers. IPS generated stronger eye-specific modulation and increased feedback connectivity to V1 during rivalry compared to replay. Ocular-layer selective activity of the LGN showed no evidence of eye-specific rivalry modulation. Experiment 3 used M and P pathway-selective visual stimuli to further investigate hierarchical neural processes of perceptual rivalry over the whole brain. Robust perceptual rivalry modulation was found in P- but not M-biased voxels in the geniculostriate pathway. IPS and ventral pulvinar showed robust perceptual modulation in rivalry and increased connectivity to the early visual cortex compared to replay.

### Interocular competition arises from lateral mutual inhibition between V1 ODCs

Horizontal connections between orientation-selective neurons are most prominent in layers 2/3 of V1 (Angelucci et al., 2017; Gilbert & Wiesel, 1983). Inhibitory large basket cells spanning ODCs were found in layer 3 of cat V1 (Buzs et al., 2001). Dichoptic cross-orientation suppression is strongest in the superficial layers of V1 in both anesthetized cats and awake monkeys (M. A. Cox et al., 2019; Sengpiel et al., 1995). According to these studies, our findings that eye-specific rivalry modulation was strongest in the superficial layers of V1 ODCs (Fig. 2f) but was weak or absent in the LGN (Fig. 5d), clearly support that local interocular competition in binocular rivalry arises from lateral mutual inhibition between V1 ODCs. The eye-specific effect in V1 was unlikely driven by pulvino-cortical input targeting the superficial layers (Shipp, 2003), since we found no eye-specific modulation in the pulvinar. Instead, our data suggest that cortico-cortical feedbacks might play roles to synchronize and resolve interocular competitions. Eye-specific rivalry dynamics was more coherent in V1 superficial and deep layers (Fig. 3c), and IPS showed stronger eye-specific modulation and connectivity to the early visual cortex during rivalry compared to replay (Fig. 4). These findings suggest that eye-specific feedback from IPS might help to synchronize and resolve local interocular competitions in V1 into a coherent perceptual representation. Our results also support that the discrepancy between single-unit and LFP/fMRI results of perceptual suppression could be due to intracortical processing and feedback modulation of synaptic input (Maier et al., 2008), which might influence the timing rather than firing rate of neuronal output.

With ocular-layer selective activity robustly resolved with 7T fMRI (Fig. 5a, Fig. S5), we found no evidence of eye-specific rivalry modulation in the stimulus region of the LGN (Fig. 5d). This finding is consistent with electrophysiological studies showing no effect of binocular rivalry on the spike rate of LGN neurons in alert monkeys (Lehky & Maunsell, 1996). However, a 3T fMRI study found significant eye-specific effect of binocular rivalry in ocular-biased voxels of the human LGN (Haynes et al., 2005). What might underlie this discrepancy? In the previous study, a larger stimulus (bilaterally presented 120-deg wedges subtending from 1.5 to 7.5 deg of eccentricity, compared to the 0.5 to 4.5 deg disc in the current study) was used to map the ocular bias of voxels from more peripheral visual field. Given that the temporal-nasal asymmetry of attention is more pronounced in the peripheral visual field (Rafal et al., 1991) and that attention can strongly modulate LGN activity (McAlonan et al., 2008; O’Connor et al., 2002), the eye-specific effect might be a result of attentional modulation due to temporal-nasal asymmetry. Consistent with this hypothesis, we also found an eye-specific suppression effect outside the ocular-bias clusters (Fig. S6), likely due to attentional suppression outside of the stimulus region (Shmuel et al., 2002; Tootell et al., 1998). Therefore, although binocular suppression exists in the primate LGN (Kacie Dougherty et al., 2021; Schroeder et al., 1990), binocular rivalry does not strongly modulate the ocular-layer selective activity.

### Parvocellular feedback to the LGN may serve as a thalamic gatekeeper of perception-related signals

Although we found no evidence of eye-specific rivalry modulation in the LGN, perceptual rivalry between chromatic and achromatic gratings strongly modulated LGN activity in its parvocellular subdivision (Fig. 6c). Modulation amplitudes were comparable between rivalry and replay, similar to the higher-order visual areas such as hV4 (Fig. 7a). Moreover, hierarchical clustering analysis revealed closer relationship between rivalry modulations in the LGN and those in high-order brain areas (Fig. 7e). Therefore, the effect of perceptual rivalry in the LGN should be a result of feedback modulation, likely through pathway-specific corticogeniculate connections (Briggs & Usrey, 2011). Since perceptual rivalry selectively modulated P but not M responses, this effect cannot be explained by attentional modulation that would influence both M and P activity (Schneider, 2011; Schneider & Kastner, 2009). With exogenous flash suppression, Wilke and colleagues found an effect of perceptual suppression on low-frequency LFP power in the LGN of alert monkeys (Wilke et al., 2009). However, this effect might be related to the disengagement of attention upon target disappearance. Another 3T fMRI study found a correlation of LGN activity with contrast perception during binocular rivalry between a pair of low and high contrast gratings (Wunderlich et al., 2005). However, this effect could also be explained by a multiplicative effect of attention on contrast responses. Our high-resolution fMRI approach also minimized the risk of contamination of LGN signals by activity from the ventral lateral pulvinar. Therefore, our results provide clear evidence that perceptual feedbacks can strongly modulate parvocellular activity of the LGN, which might serve as a gating mechanism for conscious perception-related processing at the thalamic level.

### Binocular rivalry mainly occurs in the parvocellular visual pathway

Stronger perceptual rivalry modulation in P compared to M pathway (Fig. 7a/b) provides direct neuroimaging evidence that binocular rivalry is primarily a P-pathway phenomenon. This finding is consistent with psychophysics studies showing weak rivalry suppression of M-biased visual stimuli (He et al., 2005). The dissociation of M and P pathways in binocular rivalry can be understood in terms of their distinct functional roles and neurophysiological properties. The highly sensitive and transient nature of the M pathway supports fast detection of and immediate action to potentially important events, functions that may not require consciousness or the relatively slow feedback processes. In contrast, the function of the P pathway is to encode spatial details and color information for accurate object recognition, which may require an inferential process with feedback modulation to refine information coding. Different involvement of parallel pathways in binocular rivalry could be also due to their differences in the neural circuitry of binocular interactions. Recent studies show that binocular facilitation occurs at low contrast level (M-biased) and at an early stage of visual processing, while binocular suppression occurs at high contrast level (P-biased) and at a later stage (K Dougherty et al., 2019; Mitchell et al., 2022).

### Perceptual state-related feedback from IPS help resolve visual competition in the early visual cortex

Previous studies mainly focused on the role of frontoparietal attention network in perceptual transitions of bi-stable perception (Brascamp et al., 2018). Due to technical limitations in resolving weak and possibly finescale perceptual representations in frontoparietal regions, whether frontoparietal activity represents perceptual state in bi-stable perception remains unclear. Our 7T fMRI results revealed robust perceptual state representations in IPS during binocular rivalry (Fig. 7c), and increased feedback connectivity to the visual cortex in rivalry compared to replay (Fig. 7d), suggesting that perceptual state-related feedback from IPS might play an active role to resolve visual competitions in the early visual cortex. Since we trained the state classifier based on data from the functional localizer in which subjects performed a central fixation task, the state-related activity in IPS was unlikely due to active reports. In addition, IPS represented perceptual eye dominance in binocular rivalry but not stimulus eye-of-origin in simulated replay (Fig. 4c). Since subjects were not aware of the eye-of-origin information, the role of IPS could be resolving visual competitions in the early visual cortex even without representing the content of visual consciousness. Given the critical role of IPS in attention, the current findings are also consistent with previous studies that binocular rivalry requires attention (Brascamp & Blake, 2012; Zhang et al., 2011), but not awareness of interocular conflict (Xu et al., 2016; Zou et al., 2016).

### Ventral pulvinar regulates perceptual state-related feedback connectivity across hierarchy

Using parallel pathway-selective visual stimuli, BOLD responses in both lateral (P) and medial (M) subdivisions of ventral pulvinar (Fig. 6b) subregions significantly correlated with conscious perception during binocular rivalry (Fig. 6d, Fig. 7b). The stimulus-specific perceptual modulation cannot be explained by a non-specific effect of spatial attention. This poses an advantage over the study by Wilke et al., where target disappearance was induced by flash suppression, potentially inducing a non-specific effect of attention. However, it remains possible that perceptual modulations in the ventral pulvinar could be related to feature-based attention. Similar to IPS, pulvinar showed robust perceptual modulation during rivalry and stronger connectivity to the visual cortex in rivalry compared to replay. Therefore, our findings support a critical role of ventral pulvinar in generating conscious visual perception, which could be regulating the feedback connectivity across cortical hierarchy and supporting cortical computations to resolve visual competition.

## Conclusions

The current study revealed the most complete picture so far about how binocular rivalry is resolved in the human brain. Interocular competition in binocular rivalry arises from lateral mutual inhibition between ocular dominance columns in V1 superficial layers. Feedback modulations from IPS further synchronize and resolve local competitions in the visual cortex into a coherent perceptual representation. The ventral pulvinar serves as the network hub regulating the feedback connectivity across cortical hierarchy to resolve perceptual conflicts. Finally, parvocellular feedback to the LGN might serve as a gating mechanism of perception-related signals at the thalamic level. These findings elucidate the functional roles of major brain areas involved in binocular rivalry, and their hierarchical interactions resolving visual competition to generate conscious perception. Our study also demonstrates that 7T high-resolution fMRI of fine-scale functional modules (cortical columns, laminae, and subdomains of subcortical nuclei) can help unraveling hierarchical cortical and subcortical mechanisms in humans.

## Methods and Materials

### Participants

Sixteen healthy volunteers (seven females, age 22–40 years) participated in Experiment 1. Three of them were excluded due to lack of clear OD pattern in V1, and one subject was excluded due to strong bias toward one eye. Fifteen subjects (seven females, age 22–39 years) participated in Experiment 2. Sixteen subjects (seven females, age 22–41 years) participated in Experiment 3. One subject was excluded due to response box failure, and two were excluded due to lack of significant M- or P-biased voxels in ventral pulvinar. All observers had normal or corrected-to-normal vision and gave written informed consent. Experimental protocols were approved by the Institutional Review Panel at the Institute of Biophysics, Chinese Academy of Sciences.

### Stimuli and procedures

Fig. 1a shows the stimuli and procedures for Experiment 1 and 2. For the ocular-bias localizer, to selectively activate V1 ODCs and LGN ocular layers, a high contrast checkerboard (1 deg check size, about 8-10 deg in diameter adjusted for each individual) counterphase flickering at 8 Hz was monocularly delivered to the left or right eye in alternating 24-s blocks. Two 24-s fixation blocks were included at the beginning and the end of the run. The checkerboard slowly rotated in 3.75-degree steps every second to reduce adaptation. Subjects viewed the dichoptic stimuli in the scanner with prism glasses and a cardboard divider, and reported occasional fixation-size changes by pressing a button. During binocular rivalry, red and green gratings (0.8 cycle/deg) in orthogonal orientations were dichoptically presented to the two eyes. The gratings rotated at 0.67 round/s in the same direction to prevent adaptation. The association between color and eye swapped every run so that each eye was not bound to a particular color. Subjects continuously reported whether they were seeing red, green, or a mixed percept using three buttons. In replay runs, the perception and timing of the previous rivalry run were simulated with physically alternating monocular stimulus. The transition was simulated as a blurred and alpha blended boundary rotating and swiping across the grating, gradually revealing the stimulus from the other eye. Each rivalry or replay run lasted 256 s, whereas the ocular-bias localizer run lasted 336 s. Subjects scanned 4-6 localizer runs, 4 rivalry runs, and 4 replay runs in a single session. For the bSSFP scans of S06 in Experiment 1, 6 localizer and 6 rivalry runs were collected in one session, followed by 6 localizer and 6 replay runs in another session on a different day.

In Experiment 3, achromatic and chromatic gratings were designed to preferentially activate the M or P pathway while remaining roughly balanced during rivalry. In localizer runs, the M-biased stimulus was a low contrast (30%, and for some subjects 50% if the dominant duration for the M stimulus was too short during rivalry in a pilot experiment), luminance-defined sinewave grating (0.5 cycle/deg), counterphase flickering at 10 Hz; the P-biased stimulus was a red/green isoluminant grating (0.5 cycles/deg), counterphase flickering at 4 Hz. The isoluminance of red, green and the gray in background was adjusted for each subject with a minimal-flicker procedure. The 16-s stimulus blocks were interleaved with 16-s fixation period. During stimulus blocks, the M or P stimulus was monocularly presented either to the left or to the right eye, and its orientation changed 45 degrees every 2 s (counterclockwise or clockwise in separate blocks). Each localizer run lasted 336 s. During binocular rivalry, the chromatic and achromatic gratings were dichoptically presented in orthogonal orientations, rotating at 1 round/s without flickering. Subjects pressed one of three buttons to indicate chromatic, achromatic or mixed percepts. The percept durations were re-used in replay runs to simulate the perception during binocular rivalry. Each rivalry or replay run lasted 300 s.

The mean percept duration in binocular rivalry (excluding mixed period) in Experiment 1/2/3 were 7.13 ± 2.73 s, 7.85 ± 2.33 s, 7.97 ± 2.47 s, respectively.

### MRI data acquisition

MRI data were acquired with a 7T scanner (Siemens Magnetom) using a 32-channel receive single-channel transmit head coil (NOVA medical) in the Beijing MRI Center for Brain Research (BMCBR). A bite bar was used to reduce head motion. In Experiment 1, T2*w BOLD signals from the occipital and parietal cortices were acquired with a 2D GE-EPI sequence (0.8 mm isotropic, 31 oblique-coronal slices, FOV = 128×128 mm, TE = 23 ms, TR = 2000 ms, nominal flip angle = 80°, bandwidth = 1157 Hz/pixel, partial Fourier = 6/8, GRAPPA = 3). The author C.Q. was also scanned with a 2D passband bSSFP sequence to acquire T2w BOLD signals (voxel size 0.5 × 0.5 × 1.5 mm, 2 oblique-coronal slices, FOV = 96 × 96 mm, volume acquisition time = 2400 ms for localizer and 1600 ms for rivalry/replay, TR = 5.64 ms, TE = 2.82 ms, nominal flip angle = 29° or 30°, bandwidth = 521 Hz/pix, GRAPPA = 0 for localizer and 2 for rivalry/replay). 3D passband bSSFP sequence (voxel size 0.8 × 0.8 × 0.8 mm, 10 oblique-coronal slices, FOV = 102 × 102 mm, volume acquisition time = 6 s, TR = 5.54 ms, TE = 2.77 ms, nominal flip angle = 15°, bandwidth = 471 Hz/pix, partial Fourier = 7/8 in both phase and slice direction, GRAPPA = 2) was also used in a separate localizer session to evaluate the robustness of V1 ODC pattern. The same 2D GE-EPI sequence was used, albeit with different parameters in Experiment 2 (1.2-mm isotropic voxels, 31 oblique-transversal slices, FOV = 180×180 mm, TE = 22 ms, flip angle = 78°, bandwidth = 1587 Hz/pix, GRAPPA = 2) and Experiment 3 (1.5-mm isotropic voxels, 68 oblique-transversal slices, FOV = 183×183 mm, TE = 21.6 ms, bandwidth = 1576 Hz/pix, GRAPPA = 2, multiband = 2). EPI volumes with reversed phase encoding and readout directions were also acquired for susceptibility distortion correction. For all Experiments, T1w anatomical volumes were acquired using a MP2RAGE sequence (0.7-mm isotropic voxels, FOV = 224×224 mm, 256 sagittal slices, TE = 3.05 ms, TR = 4000 ms, TI1 = 750 ms, flip angle = 4°, TI2 = 2500 ms, flip angle = 5°, bandwidth = 240 Hz/pix, partial Fourier = 7/8, GRAPPA = 3).

### MRI data analysis

#### Preprocessing

MRI data were preprocessed using AFNI (Cox, 1996), FreeSurfer (version 6.0) (Fischl, 2012), ANTs (Avants et al., 2011), and the mripy package developed in our lab (https://github.com/herrlich10/mripy). EPI volumes were corrected for slice timing, susceptibility distortion (blip-up/down method), head motion (6 parameters rigid body), and rescaled to percent signal change. To minimize the loss of spatial resolution, all spatial transformations were combined and applied in a single interpolation step (sinc method), in which the data were also up-sampled by a factor of 2 (Wang et al., 2022). The anatomical volume as well as the reconstructed surfaces were aligned to the mean of preprocessed EPI images. Slow baseline drift and the motion parameters were regressed out for both GLM and event-related average analyses. A canonical HRF (BLOCK4 in AFNI) was used for both cortical and subcortical ROIs in the GLM unless otherwise noted. For the 2D bSSFP data, motion correction was performed in-plane with three free parameters (in-plane rotation and translation) estimated from the central part of the image that was free of aliasing. Susceptibility distortion correction was safely omitted due to the very low distortion of bSSFP images.

In Experiment 3, to increase the power for detecting P-biased clusters in subcortical regions where SNR was relatively low, within-ROI smoothing was performed within anatomical masks of the LGN and pulvinar (3dBlurInMask in AFNI, FWHM = 3 mm) after motion correction. M-biased voxels were defined with unsmoothed data because they were expected to locate in thin laminae, which might easily be contaminated or even overshadowed by P-biased voxels with smoothing.

To alleviate bias induced by pial veins in laminar analysis, surface vertices with excessively high BOLD signal change (mean stimulus-driven response over 10% in localizer runs) or low EPI intensity (below 75% of the mean EPI intensity) were classified as veins and the corresponding column of voxels were excluded from further analysis. Such column-wise voxel exclusion and ROI selection (see below) were used to balance the number of voxels from different depths in the laminar analysis, aiming for a within-column comparison in activation profile in Experiment 1.

#### Surface segmentation and depth estimation

The T1w MP2RAGE anatomical volume was segmented into white matter (WM), gray matter (GM), and cerebrospinal fluid (CSF) using the automated procedure in FreeSurfer (version 6.0) with the high-resolution option (*-hires*). The results of initial segmentation were visually inspected and manually edited to eliminate dura matter, sinus, etc., ensuring correct GM boundaries. To match the up-sampled volume grid and to alleviate the vertex-missing problem during surface-to-volume projection, high density surface meshes were created by subdividing each triangular face into 4 smaller ones at the midpoint of each edge and repeated again (yielding 16 small triangles) (Polimeni et al., 2017).

The relative cortical depth for each voxel was estimated using the equivolume method (Waehnert et al., 2014) implemented in the mripy package. The neighborhood areas for a pair of nodes on the pial or smoothwm surface were approximated by summing up the area of all triangular faces surrounding the vertex on the corresponding surface mesh. A set of intermediate surfaces on specified equivolume depths were then generated according to Equation 10 in (Waehnert et al., 2014). Finally, voxel depth was computed by interpolating between two nearest equi-depth surfaces. The pial surface (WM/GM boundary) was defined to have a relative cortical depth of zero (one). The deep, middle, and superficial layers were defined to take up 30%, 35%, and 35% of the cortical thickness, respectively (Balaram et al., 2014; de Sousa et al., 2010; Liu et al., 2020). For the eye-specific pattern analysis in Fig. 3, we used 3dVol2Surf in AFNI when projecting volume data onto the surface, which employed an equidistance algorithm. According to previous studies (Liu et al., 2020; Renzo et al., 2021), equivolume and equidistance estimates of cortical depth showed only mild differences in the final results.

#### ROI definition

In Experiment 1, V1 ROIs were manually drawn on the cortical surface to select regions with a clear and roughly balanced pattern of ODCs (see Fig. S1 for the OD patterns and ROIs of all subjects). Vertices with significant ocular bias (LE-RE contrast t > 2 for LE-biased vertices, and t < -2 for RE-biased vertices) and visual response (LE+RE t > 2) were then projected to the volume space to select voxels in a column-wise manner. IPS was defined as the union of IPS0 to IPS5 in Wang15 atlas (Wang et al., 2015), whose masks were generated using the neuropythy package (Benson et al., 2018). pIPS and aIPS was defined as IPS0-2 and IPS3-5, respectively.

In Experiment 2, anatomical mask for each LGN was manually delineated in the T1w volume, and two clusters of voxels with significant ocular bias were identified for each LGN (Fig. S5 shows the ocular-biased clusters for all subjects). V1 voxels with significant ocular bias (LE-RE abs(t) > 2) and positive visual response (LE+RE > 0) were included for ROI analysis.

In Experiment 3, cortical ROIs were first defined as coarse masks on the cortical surface, then projected back to the native voxel space, and finally refined based on M-P bias and stimulus responsiveness derived from the GLM for localizer runs. Surface masks for the early visual cortices (V1, V2, V3, and hV4) were corresponding brain areas in the Benson14 atlas (Benson et al., 2018; Benson et al., 2014). The surface mask for MT+ was manually drawn on the native mesh around the main M-biased cluster (M-P t > 2, or as low as 1 for some subjects, with 3-mm FWHM surface smoothing) within TO of the Benson14 atlas. After surface-to-volume projection, only voxels with significant M-P bias and positive M or P response were kept as the M/P subdivision of these early visual areas (for V1m, V2m, V3m, MT+: M-P t > 2 and M beta > 0; for V1p, V2p, V3p, hV4: M-P t < -2 and P > 0). For the frontoparietal areas (IFJ, FEF, TPJ, pSTS, aIPS and pIPS) with weak M-P bias, surface masks were manually defined on the standard surface (std141 in AFNI/SUMA) by encircling major clusters on the group-level t map of M+P response (t > 0.5, 4-mm FWHM surface smoothing, Fig. S7). A cross-validated feature-selection procedure was then used for each individual to select the most relevant voxels that discriminated M from P stimulation (see below). For subcortical areas (LGN, vPul, and SC), group-level anatomical masks were first manually delineated on a symmetric T1w template in MNI space (Pauli et al, 2018) by an experienced experimenter (the author P.Z.), and then nonlinearly transformed to the native space of each subject using ANTs. Subcortical masks for each individual were carefully inspected and adjusted based on T1w MP2RAGE images, and were used as the anatomical reference for cluster selection and the mask for within-ROI smoothing. P-biased voxels in the LGN were selected from the dorsal nucleus (M-P t < -2 and P > 0, FWHM = 3 mm blur-in-mask to increase SNR), whereas M-biased voxels were selected from the ventral nucleus (M-P > 0 and M t > 1, no smoothing). For the ventral pulvinar, P- and M-biased voxels were selected from its lateral and medial portions, respectively, using similar 1st-level contrasts and thresholds as the LGN. The ROI for the SC was defined as visually responsive voxels in the anatomical mask (M+P t > 1).

#### Univariate and multivariate differential response

We computed the differential response between LE and RE ODCs in assessing the eye-specific modulation across cortical depths and in the DCM analysis. For univariate analysis, computing the perceptual modulation (see below) of the differential response, is equivalent to estimating the perceptual modulation separately for LE and RE ODCs and taking their average:

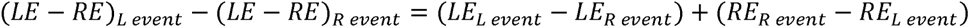

The eye-specific differential response is essentially a special linear combination of all voxels, where voxels belonging to LE ODCs are given a uniform weight of 1 while the RE voxels are all given a weight of -1. Although faithfully reflecting the mean response amplitude (e.g., across layers), this way of weight assignment may not be optimal, or even possible for higher-level areas in extracting eye-specific information. To increase sensitivity, a better set of weights can be obtained by training a linear classifier (we used linear support vector machines from the scikit-learn package with default hyper-parameters) on ocular-bias localizer data to predict which eye was stimulated on a TR-by-TR basis, and the multivariate response patterns from other conditions can then be linearly projected using the optimal weights into a 1D timeseries that reflects the distance to the decision boundary at each moment. This decoding timeseries is the multivariate differential response.

For Experiment 1, the multivariate method was used for IPS. Within atlas-defined aIPS or pIPS region, voxels with above-threshold visual response (omnibus F > 1, and L+R t > 1, but L+R beta < 5) and ocular bias (200 most biased voxels (2x up-sampled) in both ends of the L-R t distribution, with positive monocular response, e.g., LE > 0 for LE-biased voxels) based on the GLM results of ocular-bias localizer were selected as features. Results were similar across a reasonable range of thresholds. The ideal response timecourse for the localizer was created by convolving the HRF (“GAM” with default parameters in AFNI) with boxcar functions indicating LE or RE blocks, and then taking their difference. Volumes at the flat part of the block responses (absolute value of the ideal response > 0.75 * maximum) were selected for training, whereas all volumes from the rivalry/replay runs were used at test time for generating the multivariate differential response. Each sample (feature vector) was normalized to have unitary Euclidean norm before training or testing.

For Experiment 3, the multivariate method was used for frontoparietal areas and in the searchlight analysis. SVM models were trained on the localizer data to discriminate M from P stimulation. Since the spatial distribution of M vs P information across voxels might greatly differ across areas, the hyperparameters for feature selection were titrated for each ROI using grid search and cross validation. Data from rivalry and replay conditions were split into two halves with disjoint runs. The space of possible feature selection hyperparameters was sampled by a 2D grid comprising the Cartesian product of 6 levels of visual responsiveness (from no constraint to GLM omnibus F > 3, M+P t > 2, M beta > 0, P beta > 0) and 8 levels of M-P bias (from no constraint to top 1% most biased voxels (2x up-sampled) from both ends of the M-P t distribution). For each set of hyperparameters, localizer data were used to train an SVM, with which the rivalry or replay activation pattern movie was projected into 1D time series on the two halves separately. The models and the corresponding sets of features that resulted in significant perceptual modulation (see below) across subjects on the first half (the validation set) were chosen, and their results on the second half (the test set) were averaged, weighted by the effect size on the validation set. The roles of validation and test were then swapped and the average of the two test results was taken as the final result. The procedure was repeated for each ROI and each condition separately.

#### Event-related modulation

Event-related modulation, i.e., the difference in BOLD activity when subjects perceived one stimulus over the other, time-locked to button presses in either rivalry or replay condition, was estimated using several methods that generally led to similar results. To reduce the impact of HRF difference among brain areas (especially for subcortical nuclei), a model-free event-related average of BOLD signals was used in most cases. BOLD signals were averaged across voxels within each ROI (e.g., LE-biased voxels in V1 or P-biased voxels in vPul; for higher-level areas like IPS, multivariate differential response was used), linearly interpolated (0.1 s for Experiment 1/2 and 0.01 s for Experiment 3), smoothed with a 10-s hamming window (only for Experiment 3), and sorted into epochs time-aligned with button presses (LE/RE trials for Experiment 1/2 and M/P trials for Experiment 3). Trials that were shorter than 4 s or whose previous trial was shorter than 2 s were excluded. Button presses without a corresponding event in the paired rivalry or replay run were also discarded. Epochs were baseline corrected (subtracting the mean between -1 to 1 s; this was omitted for decoding timecourse in which case zero is a natural baseline) and averaged to acquire the event-related response. The modulation timecourse was obtained by subtracting responses between the L (M) and R (P) events. For eye-specific modulation, the results from LE- and RE-biased voxels were averaged. Finally, the mean value of the modulation timecourse between 4-12 s after the switch was taken as the estimated modulation amplitude. The response map of rivalry modulation was computed on the surface in a similar way but by first projecting the BOLD timeseries onto the surface. The resulting map was high-pass filtered by subtracting the smoothed version (8-mm FWHM surface smoothing). In Experiment 3, since some frontal areas exhibited large variability in the shape of the modulation timecourse, using a fixed time window for all ROIs to summarize the modulation amplitude seemed suboptimal. Thus, we determined the window using a data-driven approach based on cross-validation. For each ROI, the mean modulation timecourse across subjects for one half of the data (see previous section) was smoothed, and the interval supporting the first positive peak (within 0-15 s) was taken as the time window, within which the mean response on the other half of the data was recorded without double-dipping.

To discount the influence of sluggish BOLD signal from previous trials, we estimated the modulation amplitude for V1 laminar analysis using a GLM-based method (3dDeconvolve with CSPLINzero model in AFNI), which enjoyed the high SNR in V1. BOLD timecourse from 0 to 24 s after each perceptual switch was modeled by 11 parameters separated by 2 s, with response at 0 and 24 s fixed to be zero. Voxels for each eye and within the equivolume depth range for each layer (see above) were pooled and the averaged timeseries were fed to the model. The L and R events were modeled separately, and their results were differentiated (preferred - non-preferred) and then averaged between LE and RE ODCs to get the modulation curve. The mean response under the first positive peak was taken as the estimated modulation amplitude.

To further discount the influence of variable dominance durations across subjects in the V1-IPS correlation analysis, the multivariate differential response for each ROI was modeled by a GLM with variable length blocks (dmUBLOCK in AFNI). Besides run-wise baseline drift and head motion regressors, only one perception-related regressor was included, in which LE- and RE-dominant intervals were modeled as blocks of 1’s and -1’s before convolving with the HRF. The resulting beta value was taken as the estimated modulation amplitude.

#### Laminar analysis of eye-specific modulation

To compare the shape of laminar profiles during rivalry and replay, eye-specific modulations for each condition were normalized by dividing the sum of responses across cortical depth for each subject (Fig. 2f). Since the two conditions shared the same perception as well as laminar bias of the BOLD signal, we further calculated the rivalry/replay modulation ratio across cortical depths. To generate the continuous laminar profiles in Fig. 2e/2k, the relative cortical depth from 0 to 1 was resampled into 31 points, and the mean response at each depth was computed by averaging voxels near that depth with Gaussian weights (sigma = 0.067 for EPI and 0.1 for bSSFP). The strength of ocular dominance across layers was indexed by the amplitude of ocular modulation based on the univariate differential response in localizer runs.

#### Pattern correlation

To quantify the synchrony of eye-specific rivalry dynamics across OD columns in V1, we computed Pearson’s correlation coefficient between the moment-to-moment V1 response pattern during rivalry and replay with the ODC pattern estimated from the localizer. The preprocessed BOLD signal within each layer was projected to the surface as the instantaneous activity pattern, and the ODC pattern was computed by projecting the localizer LE-RE beta values within the gray matter to the cortical surface. The volume-to-surface projection used median map-function. Pattern correlation coefficient is less sensitive to difference in modulation amplitude across layers, because the standard deviation of the activity pattern (a spatial manifestation of the temporal modulation) is normalized in the denominator. If the ODCs for one eye become activated at slightly different times, or their modulation amplitudes vary asynchronously across the visual field, the correlation coefficient would be closer to zero on average. Thus, the width of the distribution of all r values in the TR-by-TR pattern correlation timecourse can be used as an index for the synchrony of eye-specific dynamics. The distribution width was defined as two times its standard deviation, estimated separately for each layer and condition. Since SNR also limits the maximally attainable pattern correlation, which may vary across layers, we used the distribution width in replay condition (whose response was synchronized by external stimulus drive) as a benchmark to measure the synchrony of rivalry dynamics.

In the control analysis, we tested whether the SNR difference across cortical layers alone could produce a similar laminar profile of pattern correlation. We first modeled the BOLD responses in rivalry and replay conditions and in each layer using GLMs. LE and RE dominant intervals were modeled as variable length blocks (dmUBLOCK in AFNI) in two separate regressors. After model fitting, the residual timeseries were shuffled in time independently for each vertex, which destroyed any unmodeled synchronous activity fluctuation (e.g., trial-by-trial changes in response amplitude that were synchronous across ODCs). The shuffled residuals were then added back to the fitted timeseries, and the same pattern correlation analysis was repeated for the recombined dataset.

#### Dynamic causal modeling

Effective connectivity of the fMRI data was analyzed with the DCM module of SPM12 (Version 7771). In Experiment 1, multivariate decoding timeseries from V1, V2, and IPS were used as VOI inputs. In Experiment 3, the differential timeseries between P- and M-biased voxels were used as the VOI inputs for V1 and pulvinar; the difference between the mean responses of P-biased voxels in hV4 and M-biased voxels in MT+ was used as data for the high-level visual cortex (labeled in the model as hV4); and the multivariate decoding timeseries was used as the IPS data. The timeseries of rivalry and replay conditions were concatenated and modeled together. In the eye-specific connectivity model for Experiment 1 (Fig. 4), there were two inputs: the eye-of-origin of the currently perceived stimulus (high for LE and low for RE) was defined as a driving input to V1 in both rivalry and replay conditions, and all brain areas (V1/V2/IPS) also received an additional eye-specific driving input only in rivalry. Fixed connections were defined between and within all brain areas, and the between-areas connections were allowed to be modulated by the 2nd input during binocular rivalry. Both inputs were mean-centered. A bilinear, single state, deterministic model with default parameters was used. At the first or individual level, the full DCM for each subject was estimated using all data from rivalry and replay runs (Zeidman, Jafarian, Corbin, et al., 2019). At the second or group level, we used the parametric empirical Bayes method (Friston et al., 2016; Zeidman, Jafarian, Seghier, et al., 2019) to perform Bayesian model reduction, Bayesian model average, and make inferences about the connectivity strength. Both the C matrix for the second input and the B matrix of the modulatory effect of rivalry were tested and the averaged model for explaining the commonalities across subjects was shown. Z values for the estimated parameters (e.g., changes in effective connectivity, i.e., the B matrix) were computed by dividing their expectation (Ep) with the square root of the corresponding diagonal elements in the covariance matrix (Cp). The stimulus-specific connectivity model for Experiment 3 (Fig. 7) was defined and estimated similarly.

#### Hierarchical clustering

The hierarchical clustering analysis was performed using the dendrogram function from Scipy with Euclidean distance and Ward’s linkage. The data point for each ROI was a normalized vector (with a length of one) comprising rivalry and replay modulations for all subjects. The default distance threshold of 0.7 times the maximum distance between clusters was used, and the resulting number of clusters was checked and determined from the dendrogram.

#### Statistical analysis

Statistical analyses were conducted using the Pingouin package (v0.5), JASP (v0.14), R (v4.1), and home-built Python code (for permutation and bootstrap procedures). Cluster-based permutation test (Maris et al., 2007; Nichols & Holmes, 2002) was used to test the difference in timeseries correcting for multiple comparisons (see (Ge et al., 2020) for detailed procedures). Perceptual modulations were tested against zero using one-tailed one-sample t-test and Holm correction for multiple comparisons across ROIs and conditions (or otherwise noted). Modulation differences across conditions were tested using repeated measures ANOVA followed by two-tailed paired t-test with Holm correction across ROIs (or otherwise noted). The pairwise comparisons between different layers followed by a significant ANOVA were not corrected because there were only three levels (Levin et al., 1994). The laminar profiles of perceptual modulation in rivalry and replay were normalized (so that the sum of all layers was one) before comparison. Similarly, data were normalized for each ROI by dividing the L2-norm of rivalry and replay modulations to enable comparison across ROIs in Fig. 7. The normalization would not change the test results against zero or between rivalry and replay conditions, because the modulation of each subject in each condition was divided by the same value for a given ROI.

To account for the correlation between vertices or voxels in accessing the between-session consistency of V1 ODC maps and LGN ocular-biased clusters, the observed correlation coefficients were compared with the null distribution generated by Monte Carlo simulation. The spatial auto-correlation function within the ROI was first estimated from the localizer GLM residual volumes (3dFWHMx in AFNI using the three-parameter ACF model). 10000 simulated volumes (Gaussian random noise with specified spatial smoothness) were then generated (3dClustSim in AFNI) as surrogate data, with which correlation coefficients under null hypothesis were computed to get the null distribution. Finally, the observed statistic was compared to the critical value of the null distribution for its significance.

## Acknowledgement

This study was supported by STI2030-Major Projects (2022ZD0211900, 2021ZD0204200), National Natural Science Foundation of China (31871107, 32000787, 31930053), Strategy Priority Research Program (XDB32020200) and Key Research Program of Frontier Sciences of the (KJZD-SW-L08), Youth Innovation Promotion Association project (2021089) and CAS-NWO International Cooperation Program from Chinese Academy of Science (153311KYSB20160030).

## Notes

### Competing Interest Statement

The authors have declared no competing interest.

### Summary of Updates

Title; Abstract; Add a graphical abstract; Main text; Conclusions; Discussion; Methods; Figure 1,6; Supplemental Figure S2,S4,S5,S6,S7,S8 and S9.

## References

Angelucci, A., Bijanzadeh, M., Nurminen, L., Federer, F., Merlin, S., & Bressloff, P. C. (2017). Circuits and Mechanisms for Surround Modulation in Visual Cortex. Annual Review of Neuroscience, 40, 425–451. https://doi.org/10.1146/annurev-neuro-072116-031418

Arcaro, M. J., Pinsk, M. A., & Kastner, S. (2015). The Anatomical and Functional Organization of the Human Visual Pulvinar. Journal of Neuroscience, 35(27), 9848–9871. https://doi.org/10.1523/JNEUROSCI.1575-14.2015

Avants, B. B., Tustison, N. J., Song, G., Cook, P. A., Klein, A., & Gee, J. C. (2011). A reproducible evaluation of ANTs similarity metric performance in brain image registration. Neuroimage, 54(3), 2033–2044. https://doi.org/10.1016/j.neuroimage.2010.09.025

Balaram, P., Young, N. A., & Kaas, J. H. (2014). Histological features of layers and sublayers in cortical visual areas V1 and V2 of chimpanzees, macaque monkeys, and humans. Eye Brain, 2014(6 Suppl 1), 5–18. https://doi.org/10.2147/EB.S51814

Beckett, A. J. S., Dadakova, T., Townsend, J., Huber, L., Park, S., & Feinberg, D. A. (2020). Comparison of BOLD and CBV using 3D EPI and 3D GRASE for cortical layer functional MRI at 7 T. Magnetic Resonance in Medicine, 84(6), 3128–3145. https://doi.org/10.1002/mrm.28347

Benson, N C, Jamison, K. W., Arcaro, M. J., Vu, A. T., Glasser, M. F., Coalson, T. S., Van Essen, D. C., Yacoub, E., Ugurbil, K., Winawer, J., & Kay, K. (2018). The Human Connectome Project 7 Tesla retinotopy dataset: Description and population receptive field analysis. J Vis, 18(13), 23. https://doi.org/10.1167/18.13.23

Benson, Noah C., Butt, O. H., Brainard, D. H., & Aguirre, G. K. (2014). Correction of Distortion in Flattened Representations of the Cortical Surface Allows Prediction of V1-V3 Functional Organization from Anatomy. PLoS Computational Biology, 10(3). https://doi.org/10.1371/journal.pcbi.1003538

Blake, R. (1989). A neural theory of binocular rivalry. Psychol Rev, 96(1), 145–167. http://www.ncbi.nlm.nih.gov/pubmed/2648445

Blake, R, & Logothetis, N. K. (2002). Visual competition. Nature Reviews Neuroscience, 3(1), 13–23. https://doi.org/Doi 10.1038/Nrn701

Blake, Randolph, Brascamp, J., & Heeger, D. J. (2014). Can binocular rivalry reveal neural correlates of consciousness? Philosophical Transactions of the Royal Society B-Biological Sciences, 369(1641). https://doi.org/ARTN 2013021110.1098/rstb.2013.0211

Brascamp, J. W., & Blake, R. (2012). Inattention abolishes binocular rivalry: perceptual evidence. Psychol Sci, 23(10), 1159–1167. https://doi.org/10.1177/0956797612440100

Brascamp, J, Blake, R., & Knapen, T. (2015). Negligible fronto-parietal BOLD activity accompanying unreportable switches in bistable perception. Nat Neurosci, 18(11), 1672–1678. https://doi.org/10.1038/nn.4130

Brascamp, Jan, Sterzer, P., Blake, R., & Knapen, T. (2018). Multistable Perception and the Role of the Frontoparietal Cortex in Perceptual Inference. Annual Review of Psychology, 69(1), 77–103. https://doi.org/10.1146/annurev-psych-010417-085944

Bridge, H., Leopold, D. A., & Bourne, J. A. (2016). Adaptive Pulvinar Circuitry Supports Visual Cognition. Trends in Cognitive Sciences, 20(2), 146–157. https://doi.org/10.1016/j.tics.2015.10.003

Briggs, F., & Usrey, W. M. (2011). Corticogeniculate feedback and visual processing in the primate. J Physiol, 589(Pt 1), 33–40. https://doi.org/10.1113/jphysiol.2010.193599

Burkhalter, A., & Van Essen, D. C. (1986). Processing of color, form and disparity information in visual areas VP and V2 of ventral extrastriate cortex in the macaque monkey. Journal of Neuroscience, 6(8), 2327–2351. https://doi.org/10.1523/jneurosci.06-08-02327.1986

Buzs, P., Eysel, U. T., Adorjn, P., & Kisvrday, Z. F. (2001). Axonal topography of cortical basket cells in relation to orientation, direction, and ocular dominance maps. Journal of Comparative Neurology, 437(3), 259–285. https://doi.org/10.1002/cne.1282

Carmel, D., Walsh, V., Lavie, N., & Rees, G. (2010). Right parietal TMS shortens dominance durations in binocular rivalry. Curr Biol, 20(18), R799–800. https://doi.org/10.1016/j.cub.2010.07.036

Cox, M. A., Dougherty, K., Westerberg, J. A., Schall, M. S., & Maier, A. (2019). Temporal dynamics of binocular integration in primary visual cortex. 19, 1–21.

Cox, R. W. (1996). AFNI: software for analysis and visualization of functional magnetic resonance neuroimages. Comput Biomed Res, 29(3), 162–173. https://doi.org/10.1006/cbmr.1996.0014

Crick, F. (1996). Visual perception: rivalry and consciousness. Nature.

Dayan, P. (1998). A hierarchical model of binocular rivalry. Neural Comput, 10(5), 1119–1135. http://www.ncbi.nlm.nih.gov/pubmed/9654769

de Hollander, G., van der Zwaag, W., Qian, C., Zhang, P., & Knapen, T. (2021). Ultra-high field fMRI reveals origins of feedforward and feedback activity within laminae of human ocular dominance columns. NeuroImage, 228(December 2020), 117683. https://doi.org/10.1016/j.neuroimage.2020.117683

de Jong, M. C., Vansteensel, M. J., van Ee, R., Leijten, F. S. S., Ramsey, N. F., Dijkerman, H. C., Dumoulin, S. O., & Knapen, T. (2020). Intracranial Recordings Reveal Unique Shape and Timing of Responses in Human Visual Cortex during Illusory Visual Events. Current Biology, 30(16), 3089–3100.e4. https://doi.org/10.1016/j.cub.2020.05.082

de Sousa, A. A., Sherwood, C. C., Schleicher, A., Amunts, K., MacLeod, C. E., Hof, P. R., & Zilles, K. (2010). Comparative Cytoarchitectural Analyses of Striate and Extrastriate Areas in Hominoids. Cerebral Cortex, 20(4), 966–981. https://doi.org/10.1093/cercor/bhp158

Derrington, A. M., & Lennie, P. (1984). Spatial and temporal contrast sensitivities of neurones in lateral geniculate nucleus of macaque. J Physiol, 357, 219–240. http://www.ncbi.nlm.nih.gov/entrez/query.fcgi?cmd=Retrieve&db=PubMed&dopt=Citation&list_uids=6512690

Dougherty, K, Cox, M. A., Westerberg, J. A., & Maier, A. (2019). Binocular Modulation of Monocular V1 Neurons. Curr Biol, 29(3), 381–391 e4. https://doi.org/10.1016/j.cub.2018.12.004

Dougherty, Kacie, Carlson, B. M., Cox, M. A., Westerberg, J. A., Zinke, W., Schmid, M. C., Martin, P. R., & Maier, A. (2021). Binocular suppression in the macaque lateral geniculate nucleus reveals early competitive interactions between the eyes. ENeuro, 8(2), 1–12. https://doi.org/10.1523/ENEURO.0364-20.2020

Dougherty, Kacie, Schmid, M. C., & Maier, A. (2018). Binocular response modulation in the lateral geniculate nucleus. J Comp Neurol, 527(3), 522–534. https://doi.org/10.1002/cne.24417

Felleman, D. J., & Van Essen, D. C. (1991). Distributed hierarchical processing in the primate cerebral cortex. Cereb Cortex, 1(1), 1–47. https://www.ncbi.nlm.nih.gov/pubmed/1822724

Fischl, B. (2012). FreeSurfer. Neuroimage, 62(2), 774–781.

Friston, K. J., Litvak, V., Oswal, A., Razi, A., Stephan, K. E., van Wijk, B. C. M., Ziegler, G., & Zeidman, P. (2016). Bayesian model reduction and empirical Bayes for group (DCM) studies. Neuroimage, 128, 413–431. https://doi.org/10.1016/j.neuroimage.2015.11.015

Ge, Y., Zhou, H., Qian, C., Zhang, P., Wang, L., & He, S. (2020). Adaptation to feedback representation of illusory orientation produced from flash grab effect. Nature Communications, 11(1). https://doi.org/10.1038/s41467-020-17786-1

Gilbert, C. D., & Wiesel, T. N. (1983). Clustered intrinsic connections in cat visual cortex. Journal of Neuroscience, 3(5), 1116–1133. https://doi.org/10.1523/jneurosci.03-05-01116.1983

Guillery, R. W., & Colonnier, M. (1970). Synaptic patterns in the dorsal lateral geniculate nucleus of the monkey. Zeitschrift Für Zellforschung Und Mikroskopische Anatomie, 103(1), 90–108. https://doi.org/10.1007/BF00335403

Haynes, J. D., Deichmann, R., & Rees, G. (2005). Eye-specific effects of binocular rivalry in the human lateral geniculate nucleus. Nature, 438(7067), 496–499. https://doi.org/nature04169 [pii]10.1038/nature04169

He, S., Carlson, T., & Chen, X. (2005). Parallel Pathways and Temporal Dynamics in Binocular Rivalry. Binocular Rivalry.

Jaramillo, J., Mejias, J. F., & Wang, X. J. (2019). Engagement of Pulvino-cortical Feedforward and Feedback Pathways in Cognitive Computations. Neuron, 101(2), 321-+. https://doi.org/10.1016/j.neuron.2018.11.023

Kaas, J. H., & Lyon, D. C. (2007). Pulvinar contributions to the dorsal and ventral streams of visual processing in primates. In Brain Research Reviews (Vol. 55, Issue 2 SPEC. ISS., pp. 285–296). https://doi.org/10.1016/j.brainresrev.2007.02.008

Kanai, R., Carmel, D., Bahrami, B., & Rees, G. (2011). Structural and functional fractionation of right superior parietal cortex in bistable perception. Curr Biol, 21(3), R106–7. https://doi.org/10.1016/j.cub.2010.12.009

Kapoor, V., Dwarakanath, A., Safavi, S., Werner, J., Besserve, M., Panagiotaropoulos, T. I., & Logothetis, N. K. (2022). Decoding internally generated transitions of conscious contents in the prefrontal cortex without subjective reports. Nature Communications, 13(1). https://doi.org/10.1038/s41467-022-28897-2

Kovács, I., Papathomas, T. V., Yang, M., & Fehér, Á. (1996). When the brain changes its mind: Interocular grouping during binocular rivalry. Proceedings of the National Academy of Sciences of the United States of America, 93(26), 15508–15511. https://doi.org/10.1073/pnas.93.26.15508

Lehky, S. R., & Maunsell, J. H. R. (1996). No binocular rivalry in the LGN of alert macaque monkeys. Vision Res, 36(9), 1225–1234. https://doi.org/Doi 10.1016/0042-6989(95)00232-4

Leopold, D. A., & Logothetis, N. K. (1996). Activity changes in early visual cortex reflect monkeys’ percepts during binocular rivalry. Nature, 379(6565), 549–553. https://doi.org/10.1038/379549a0

Levin, J. R., Serlin, R. C., & Seaman, M. A. (1994). A controlled, powerful multiple-comparison strategy for several situations. Psychological Bulletin, 115(1), 153.

Li, H. H., Rankin, J., Rinzel, J., Carrasco, M., & Heeger, D. J. (2017). Attention model of binocular rivalry. Proc Natl Acad Sci U S A, 114(30), E6192–E6201. https://doi.org/10.1073/pnas.1620475114

Liu, C., Guo, F., Qian, C., Zhang, Z., Sun, K., Wang, D. J. J., He, S., & Zhang, P. (2020). Layer-dependent multiplicative effects of spatial attention on contrast responses in human early visual cortex. Progress in Neurobiology, 207, 101897. https://doi.org/https://doi.org/10.1016/j.pneurobio.2020.101897

Logothetis, N. K., & Wandell, B. A. (2004). Interpreting the BOLD signal. Annu Rev Physiol, 66, 735–769. https://doi.org/10.1146/annurev.physiol.66.082602.092845

Lumer, E. D., Friston, K. J., & Rees, G. (1998). Neural correlates of perceptual rivalry in the human brain. Science, 280(5371), 1930–1934. http://www.ncbi.nlm.nih.gov/pubmed/9632390

Maier, A., Wilke, M., Aura, C., Zhu, C., Ye, F. Q., & Leopold, D. A. (2008). Divergence of fMRI and neural signals in V1 during perceptual suppression in the awake monkey. Nat Neurosci, 11(10), 1193–1200. https://doi.org/10.1038/nn.2173

Maris, E., Schoffelen, J.-M., & Fries, P. (2007). Nonparametric statistical testing of coherence differences. Journal of Neuroscience Methods, 163(1), 161–175.

Mashour, G. A., Roelfsema, P., Changeux, J. P., & Dehaene, S. (2020). Conscious Processing and the Global Neuronal Workspace Hypothesis. Neuron, 105(5), 776–798. https://doi.org/10.1016/j.neuron.2020.01.026

Maunsell, J. H. R., & Van Essen, D. C. (1983). Functional properties of neurons in middle temporal visual area of the macaque monkey. II. Binocular interactions and sensitivity to binocular disparity. Journal of Neurophysiology, 49(5), 1148–1167. https://doi.org/10.1152/jn.1983.49.5.1148

McAlonan, K., Cavanaugh, J., & Wurtz, R. H. (2008). Guarding the gateway to cortex with attention in visual thalamus. Nature, 456(7220), 391–394. https://doi.org/10.1038/nature07382

Mitchell, B. A., Dougherty, K., Westerberg, J. A., Carlson, B. M., Daumail, L., Maier, A., & Cox, M. A. (2022). Stimulating both eyes with matching stimuli enhances V1 responses. IScience, 25(5). https://doi.org/10.1016/j.isci.2022.104182

Mo, C., Lu, J., Shi, C., & Fang, F. (2022). Neural representations of competing stimuli along the dorsal and ventral visual pathways during binocular rivalry. *Cerebral Cortex*, bhac238. https://doi.org/10.1093/cercor/bhac238

Moon, C. H., Fukuda, M., Park, S. H., & Kim, S. G. (2007). Neural interpretation of blood oxygenation level-dependent fMRI maps at submillimeter columnar resolution. Journal of Neuroscience, 27(26), 6892–6902. https://doi.org/10.1523/Jneurosci.0445-07.2007

Myerson, J., Miezin, F., & Allman, J. (1981). Binocular rivalry in macaque monkeys and humans: a comparative study in perception. *Behav*. Anal. Lett, 1, 149–159.

Nichols, T. E., & Holmes, A. P. (2002). Nonparametric permutation tests for functional neuroimaging: A primer with examples. Human Brain Mapping, 15(1), 1–25. https://doi.org/DOI 10.1002/hbm.1058

O’Connor, D. H., Fukui, M. M., Pinsk, M. A., & Kastner, S. (2002). Attention modulates responses in the human lateral geniculate nucleus. Nat Neurosci, 5(11), 1203–1209. http://www.ncbi.nlm.nih.gov/entrez/query.fcgi?cmd=Retrieve&db=PubMed&dopt=Citation&list_uids=12379861

O’Hashi, K., Fekete, T., Deneux, T., Hildesheim, R., van Leeuwen, C., & Grinvald, A. (2018). Interhemispheric Synchrony of Spontaneous Cortical States at the Cortical Column Level. Cerebral Cortex, 28(5), 1794–1807. https://doi.org/10.1093/cercor/bhx090

Olman, C. A., Harel, N., Feinberg, D. A., He, S., Zhang, P., Ugurbil, K., & Yacoub, E. (2012). Layer-specific fMRI reflects different neuronal computations at different depths in human V1. PLoS One, 7(3), e32536. https://doi.org/10.1371/journal.pone.0032536

Omer, D. B., Fekete, T., Ulchin, Y., Hildesheim, R., & Grinvald, A. (2018). Dynamic Patterns of Spontaneous Ongoing Activity in the Visual Cortex of Anesthetized and Awake Monkeys are Different. Cereb Cortex. https://doi.org/10.1093/cercor/bhy099

Polimeni, J. R., Renvall, V., Zaretskaya, N., & Fischl, B. (2017). Analysis strategies for high-resolution UHF-fMRI data. Neuroimage. https://doi.org/10.1016/j.neuroimage.2017.04.053

Qian, Y., Zou, J., Zhang, Z., An, J., Zuo, Z., Zhuo, Y., Wang, D. J. J., & Zhang, P. (2020). Robust functional mapping of layer-selective responses in human lateral geniculate nucleus with high-resolution 7T fMRI. Proceedings of the Royal Society B: Biological Sciences, 287(1925). https://doi.org/10.1098/rspb.2020.0245

Rafal, R., Henik, A., & Smith, J. (1991). Extrageniculate contributions to reflex visual orienting in normal humans: a temporal hemifield advantage. J Cogn Neurosci, 3(4), 322–328. https://doi.org/10.1162/jocn.1991.3.4.322

Renzo, L., Poser, B. A., Bandettini, P. A., Arora, K., Wagstyl, K., Cho, S., Goense, J., Nothnagel, N., Tyler, A., Hurk, J. Van Den, Müller, A. K., Reynolds, R. C., Glen, D. R., Goebel, R., & Faruk, O. (2021). NeuroImage LayNii : A software suite for layer-fMRI. NeuroImage, 237(February), 118091. https://doi.org/10.1016/j.neuroimage.2021.118091

Saalmann, Y. B., Pinsk, M. A., Wang, L., Li, X., & Kastner, S. (2012). The Pulvinar Regulates Information Transmission Between Cortical Areas Based on Attention Demands. Science, 337(6095), 753–756. https://doi.org/10.1126/science.1223082

Scheffler, K., Heule, R., M, G. B.-Y., Kardatzki, B., & Lohmann, G. (2018). The BOLD sensitivity of rapid steady-state sequences. Magn Reson Med. https://doi.org/10.1002/mrm.27585

Schneider, K. A. (2011). Subcortical mechanisms of feature-based attention. Journal of Neuroscience, 31(23), 8643–8653. https://doi.org/10.1523/JNEUROSCI.6274-10.2011

Schneider, K. A., & Kastner, S. (2009). Effects of sustained spatial attention in the human lateral geniculate nucleus and superior colliculus. Journal of Neuroscience, 29(6), 1784–1795. http://www.ncbi.nlm.nih.gov/pubmed/19211885

Schroeder, C. E., Tenke, C. E., Arezzo, J. C., & Vaughan, H. G. (1990). Binocularity in the Lateral Geniculate-Nucleus of the Alert Macaque. Brain Research, 521(1–2), 303–310. https://doi.org/Doi 10.1016/0006-8993(90)91556-V

Schwarzkopf, D. S., Schindler, A., & Rees, G. (2010). Knowing with which eye we see: Utrocular discrimination and eye-specific signals in human visual cortex. PLoS ONE, 5(10). https://doi.org/10.1371/journal.pone.0013775

Sengpiel, F., Blakemore, C., & Harrad, R. (1995). Interocular suppression in the primary visual cortex: a possible neural basis of binocular rivalry. Vision Res, 35(2), 179–195. https://doi.org/0042-6989(94)00125-6 [pii]

Shipp, S. (2003). The functional logic of cortico-pulvinar connections. Philos Trans R Soc Lond B Biol Sci, 358(1438), 1605–1624. https://doi.org/10.1098/rstb.2002.1213

Shmuel, A., Yacoub, E., Pfeuffer, J., Van de Moortele, P. F., Adriany, G., Hu, X., & Ugurbil, K. (2002). Sustained negative BOLD, blood flow and oxygen consumption response and its coupling to the positive response in the human brain. Neuron, 36(6), 1195–1210. https://www.ncbi.nlm.nih.gov/pubmed/12495632

Tong, F., & Engel, S. A. (2001). Interocular rivalry revealed in the human cortical blind-spot representation. Nature, 411(6834), 195–199. https://doi.org/10.1038/3507558335075583 [pii]

Tong, F., Meng, M., & Blake, R. (2006). Neural bases of binocular rivalry. Trends Cogn Sci, 10(11), 502–511. https://doi.org/S1364-6613(06)00240-3 [pii]10.1016/j.tics.2006.09.003

Tononi, G., Boly, M., Massimini, M., & Koch, C. (2016). Integrated information theory: From consciousness to its physical substrate. Nature Reviews Neuroscience, 17(7), 450–461. https://doi.org/10.1038/nrn.2016.44

Tootell, R. B. H., Hadjikhani, N., Hall, E. K., Marrett, S., Vanduffel, W., Vaughan, J. T., & Dale, A. M. (1998). The retinotopy of visual spatial attention. Neuron, 21(6), 1409–1422.

Uludag, K., & Havlicek, M. (2021). Determining laminar neuronal activity from BOLD fMRI using a generative model. Progress in Neurobiology, 207(March), 102055. https://doi.org/https://doi.org/10.1016/j.pneurobio.2021.102055

Waehnert, M. D., Dinse, J., Weiss, M., Streicher, M. N., Waehnert, P., Geyer, S., Turner, R., & Bazin, P. L. (2014). Anatomically motivated modeling of cortical laminae. Neuroimage, 93 *Pt* *2*, 210–220. https://doi.org/10.1016/j.neuroimage.2013.03.078

Wang, J., Nasr, S., Roe, A. W., & Polimeni, J. R. (2022). Critical factors in achieving fine-scale functional MRI : Removing sources of inadvertent spatial smoothing . *Human Brain Mapping*, November 2021, 1–21. https://doi.org/10.1002/hbm.25867

Wang, L., Mruczek, R. E. B., Arcaro, M. J., & Kastner, S. (2015). Probabilistic maps of visual topography in human cortex. Cerebral Cortex, 25(10), 3911–3931. https://doi.org/10.1093/cercor/bhu277

Wiesel, T. N., & Hubel, D. H. (1966). Spatial and chromatic interactions in the lateral geniculate body of the rhesus monkey. J Neurophysiol, 29(6), 1115–1156. http://www.ncbi.nlm.nih.gov/pubmed/4961644

Wilke, M., Mueller, K. M., & Leopold, D. A. (2009). Neural activity in the visual thalamus reflects perceptual suppression. Proc Natl Acad Sci U S A, 106(23), 9465–9470. https://doi.org/10.1073/pnas.0900714106

Wilson, H. R. (2003). Computational evidence for a rivalry hierarchy in vision. Proc Natl Acad Sci U S A, 100(24), 14499–14503. https://doi.org/10.1073/pnas.2333622100

Wunderlich, K., Schneider, K. A., & Kastner, S. (2005). Neural correlates of binocular rivalry in the human lateral geniculate nucleus. Nat Neurosci, 8(11), 1595–1602. https://doi.org/nn1554 [pii]10.1038/nn1554

Xu, H., Han, C., Chen, M., Li, P., Zhu, S., Fang, Y., Hu, J., Ma, H., & Lu, H. D. (2016). Rivalry-Like Neural Activity in Primary Visual Cortex in Anesthetized Monkeys. Journal of Neuroscience, 36(11), 3231–3242. https://doi.org/10.1523/JNEUROSCI.3660-15.2016

Zaretskaya, N., Bause, J., Polimeni, J. R., Grassi, P. R., Scheffler, K., & Bartels, A. (2020). Eye-selective fMRI activity in human primary visual cortex: Comparison between 3 T and 9.4 T, and effects across cortical depth. NeuroImage, 220(October 2019). https://doi.org/10.1016/j.neuroimage.2020.117078

Zaretskaya, N., Thielscher, A., Logothetis, N. K., & Bartels, A. (2010). Disrupting parietal function prolongs dominance durations in binocular rivalry. Curr Biol, 20(23), 2106–2111. https://doi.org/10.1016/j.cub.2010.10.046

Zeidman, P., Jafarian, A., Corbin, N., Seghier, M. L., Razi, A., Price, C. J., & Friston, K. J. (2019). A guide to group effective connectivity analysis, part 1: First level analysis with DCM for fMRI. Neuroimage, 200, 174–190. https://doi.org/10.1016/j.neuroimage.2019.06.031

Zeidman, P., Jafarian, A., Seghier, M. L., Litvak, V., Cagnan, H., Price, C. J., & Friston, K. J. (2019). A guide to group effective connectivity analysis, part 2: Second level analysis with PEB. Neuroimage, 200, 12–25. https://doi.org/10.1016/j.neuroimage.2019.06.032

Zhang, P., Jamison, K., Engel, S., He, B., & He, S. (2011). Binocular rivalry requires visual attention. Neuron, 71(2), 362–369. https://doi.org/10.1016/j.neuron.2011.05.035 S0896-6273(11)00492-2 [pii]

Zhang, P., Jiang, Y., & He, S. (2012). Voluntary attention modulates processing of eye-specific visual information. Psychol Sci, 23(3), 254–260. https://doi.org/10.1177/09567976114242890956797611424289 [pii]

Zhou, H., Schafer, R. J., & Desimone, R. (2016). Pulvinar-Cortex Interactions in Vision and Attention. Neuron, 89(1), 209–220. https://doi.org/10.1016/j.neuron.2015.11.034

Zou, J., He, S., & Zhang, P. (2016). Binocular rivalry from invisible patterns. Proceedings of the National Academy of Sciences of the United States of America, 113(30). https://doi.org/10.1073/pnas.1604816113

