## Supplemental Figures for "Hierarchical and fine-scale mechanisms of binocular rivalry for conscious perception"

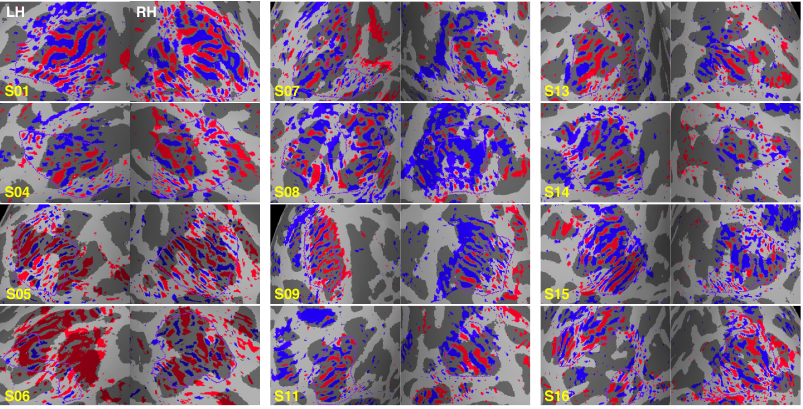

**Figure S1.** **V1 ODC patterns for all participants in Experiment 1.** Maps were thresholded at LE-RE p < 0.05 uncorrected. Solid purple lines indicate the boundary of manually defined surface ROIs.

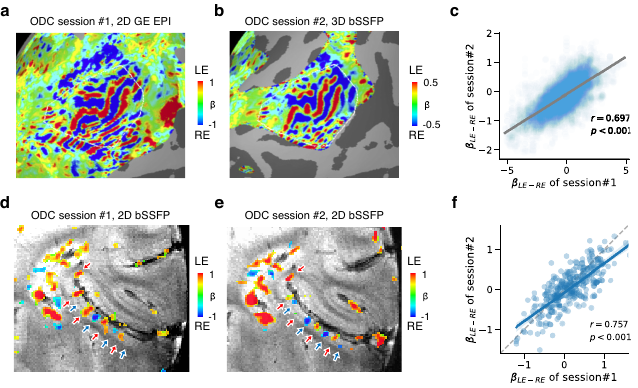

**Figure S2.** **ODC patterns in V1 were highly reproducible across different fMRI sessions.** **(a-c)** ODC patterns of S01 acquired with GE-EPI and 3D bSSFP sequences in different days. The correlation in (c) was computed within the region encircled by the white line. The p-value was obtained by a Monte Carlo test to account for the spatial dependency between vertices or voxels. **(d-f)** ODC patterns of S06 acquired with 2D bSSFP sequence in two different days. The red (blue) arrows near LE (RE) ODCs are guides for visual comparisons.

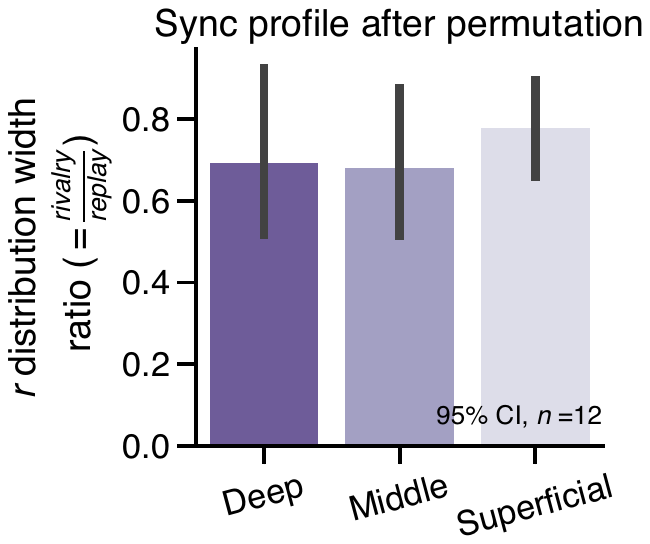

**Figure S3. Pattern synchronization in different V1 depths after temporal permutation.** The ratio between rivalry and replay for the width of distribution of moment-to-moment pattern correlation coefficients showed no significant difference across layers. Bayes factor analysis supported that the difference between the deep and middle layer was more likely to be observed under the null hypothesis that there was no difference (BF_01_=3.257, n=12).

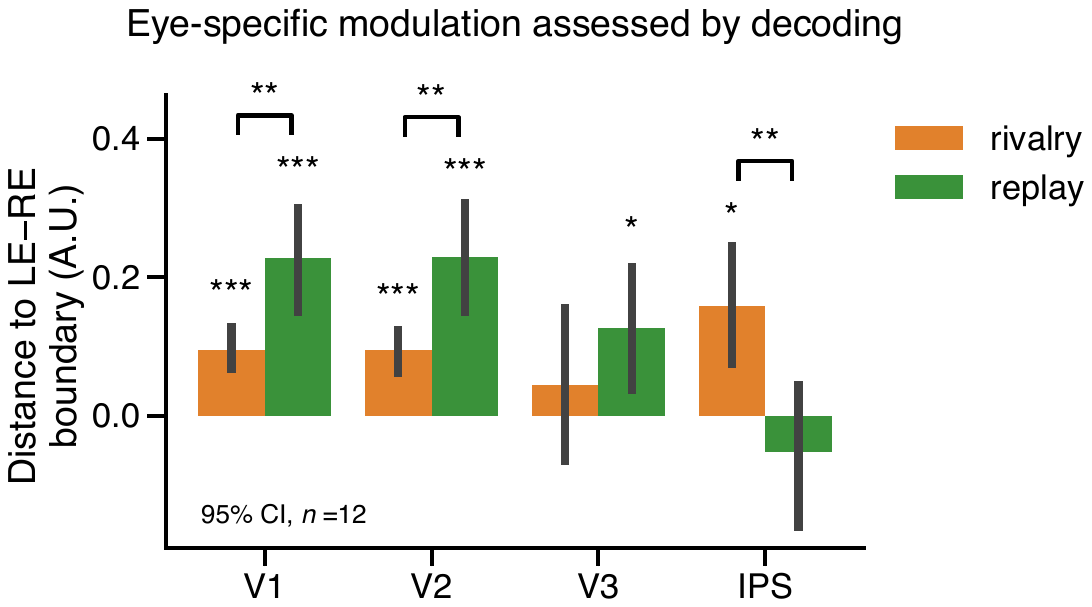

**Figure S4. Eye-specific modulations in the early visual cortex and IPS.** The modulation amplitude was calculated as the multivariate distance to the LE-RE decision boundary, normalized (divided) by the L2-norm of the data vector (rivalry and replay modulation of all subjects).

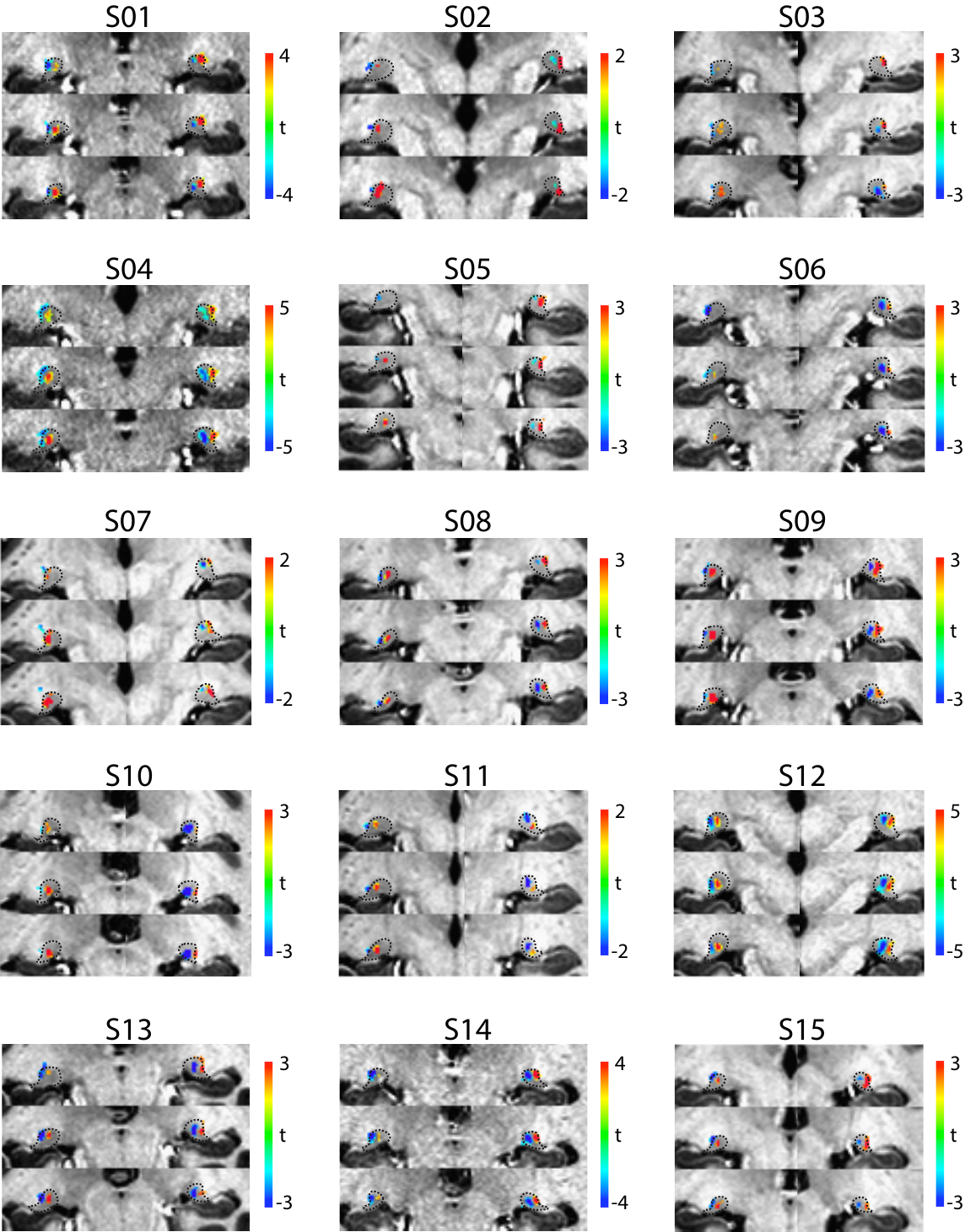

**Figure S5.** **Ocular-biased clusters of the LGNs for all participants in Experiment 2.** Colorbars encode t values for the LE-RE contrast in the GLM for ocular-bias localizers. The dotted lines mark the anatomical boundary of the LGN in three coronal slices.

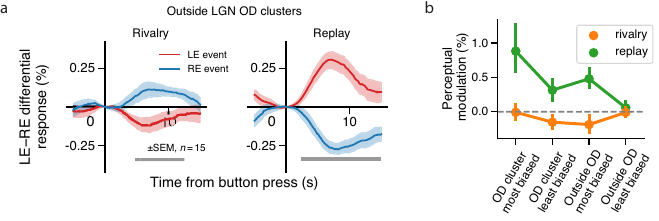

**Figure S6.** **(a)** Eye-specific modulation in LGN voxels outside the ocular-biased clusters. The differential response between LE- and RE-biased voxels (top 50 voxels in both ends of the LE-RE t distribution with omnibus F > 1 and LE+RE t > 1) was negatively modulated during binocular rivalry, likely due to attentional suppression outside the stimulus region. **(b)** Eye-specific modulation in the LGN as a function of ocular bias from inside to outside of the ocular-biased clusters. There was no eye-specific rivalry modulation in voxels with the strongest ocular-bias (top 50 voxels in both ends of the LE-RE t distribution with omnibus F > 3 and LE+RE t > 2) inside the ocular-based clusters (BF_01_ = 3.759, n = 15). Error bars indicate 95% confidence intervals.

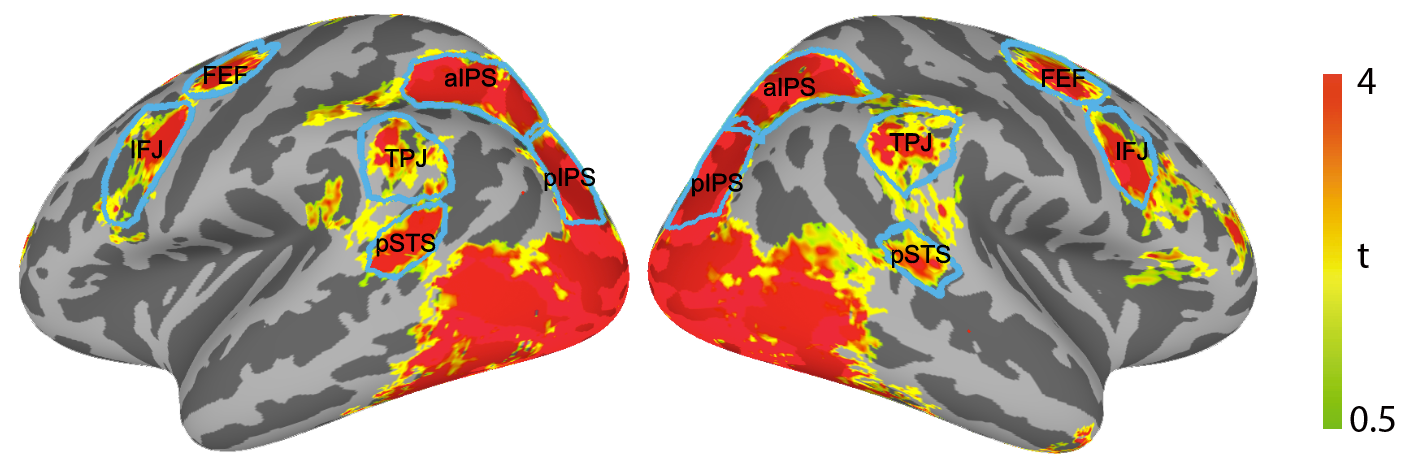

**Figure S7. ROI definitions of frontoparietal areas.** Frontoparietal ROIs were defined based on the responsiveness to the visual stimulus used in Experiment 3 (group-level M+P t map across subjects in the functional localizers, thresholded at M+P t > 0.5).

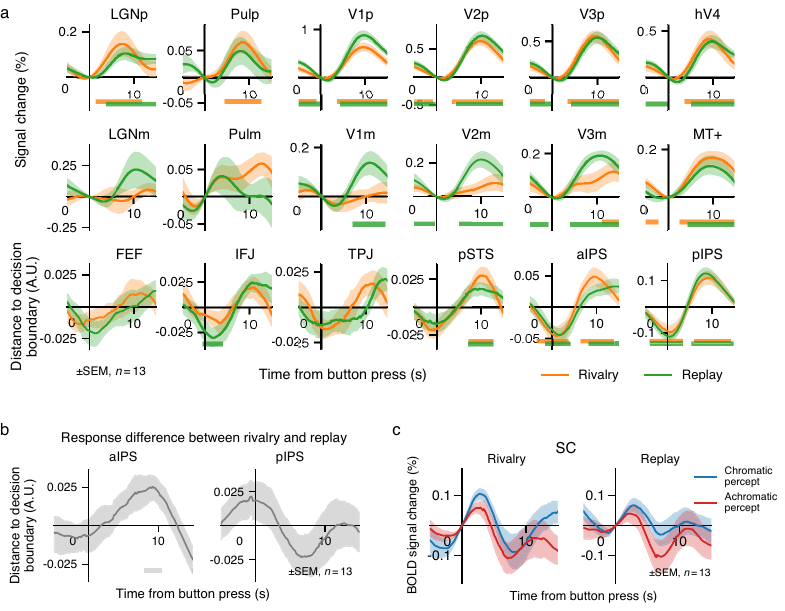

**Figure S8.** **(a)** Event-related timecourses of rivalry (orange) and replay (green) modulations for all ROIs in Experiment 3. For M- and P-biased visual areas, the differential timecourse was calculated as the difference in BOLD responses between preferred and non-preferred perceptual events. For frontoparietal areas, the differential timecourse was the difference in decoding timecourse (distance to M-P decision boundary) between the M and P perceptual events. Horizontal bars denote time periods significantly different from 0 (*p* < 0.05 corrected). **(b)** The differential timecourses between rivalry and replay conditions in aIPS and pIPS. The light gray bar denotes the time period significantly different from 0 (*p* < 0.05 uncorrected). **(c)** Event-related modulation in the SC time-locked to the chromatic (P) and achromatic (M) perceptual events. Shaded bands represent standard error of the mean (SEM) in all panels.

**Figure S9. (a)** Group-level t maps for the localizer M-P contrast. Red (blue) vertices showed significantly stronger activation to the M(P)-biased stimulus. **(b)** Group-level searchlight results based on multivariate differential response calculated as the distance to the SVM decision boundary. The value of each vertex represents the *t* statistic of event-related perceptual modulation across subjects. The surface-based searchlight was performed within disc patches of 6 mm radius centered on each vertex on the unsmoothed surface. For rivalry and replay conditions, localizer data were used as training dataset. For the localizer map, leave-one-run-out cross-validation was used. Only clusters over 100 vertices were shown. **(c)** Group-level t maps of searchlight results in the pulvinar. Before group-level analysis, individual maps were spatially smoothed within the pulvinar with a 3-mm FWHM filter. All maps were thresholded at *p* < 0.05 uncorrected.
